# Engineering of human mini-bones for the standardized modeling of healthy hematopoiesis, leukemia and solid tumor metastasis

**DOI:** 10.1101/2021.09.11.459806

**Authors:** Ani Grigoryan, Dimitra Zacharaki, Alexander Balhuizen, Christophe RM Côme, Anne-Katrine Frank, Alejandro Garcia Garcia, Kristina Aaltonen, Adriana Mañas, Javanshir Esfandyari, Nasim Kalantari, Pontus Kjellman, Sujeethkumar Prithiviraj, Emelie Englund, Chris D Madsen, Bo Porse, Daniel Bexell, Paul E Bourgine

## Abstract

The bone marrow microenvironment provides indispensable factors to sustain blood production throughout life. It is also a hotspot for the progression of hematologic disorders and the most frequent site of solid tumor metastasis. Pre-clinical research relies on xenograft mouse models, precluding the human-specific functional interactions of stem cells with their bone marrow microenvironment. Human mesenchymal cells can be exploited for the *in vivo* engineering of humanized ossicles (hOss). Those mini-bones provide a human niche conferring engraftment of human healthy and malignant blood samples, yet suffering from major reproducibility issue. Here, we report the standardized generation of hOss by developmental priming of a custom-designed human mesenchymal cell line. We demonstrate superior engraftment of cord blood hematopoietic cells and primary acute myeloid leukemia samples, but also validate our hOss as metastatic site for breast cancer cells. Finally, we report the first engraftment of neuroblastoma patient-derived xenograft cells in a humanized model, recapitulating clinically reported osteolytic lesions. Collectively, our hOss constitute a powerful standardized and malleable platform to model normal hematopoiesis, leukemia and solid tumor metastasis.

## Introduction

Bone is a multifaceted organ ensuring essential functions. During development, mesenchymal stromal cells (MSCs) orchestrate the bone formation and ultimately persist in the bone marrow (BM) compartment ^1^. MSCs ensure the production of non-hematopoietic cells including osteoblasts, adipocytes, stromal and vessel-associated cells ^2–4^. Those cells are pivotal entities constituting the mesenchymal fraction of the so-called BM niche and tightly regulating the self-renewing and differentiation activities of hematopoietic stem cells (HSCs) for the production of all blood lineages ^5–8^. Increasing evidence is being compiled on the crucial role of mesenchymal populations in the regulation of HSC functions via both secreted cues and cell-to-cell interactions ^9, 10^. Conversely, alterations in the BM niche can disrupt tissue homeostasis and induce malignant transformation ^11, 12^. Several studies evidenced the impact of mesenchymal dysfunction in leukemia emergence, but also their capacity to further promote disease development ^13^. The BM microenvironment is also a privileged ground for a large number of metastatic solid tumors ^14–17^, where cancer stem cells can engraft but also invigorate before provoking multi-organ secondary metastases ^18^. Taken together, the BM niche and the associated mesenchymal cell populations are closely involved in the development of leukemia and solid tumors. Still, the mechanisms associated with these processes are poorly understood, predominantly due to the lack of systems conferring the opportunity to compile human-specific findings.

Xenograft mouse models can be considered as the gold standard for the study of a wide variety of human hematopoietic and solid tumor malignancies. However, several primary patient materials still poorly engraft in mice ^19, 20^, challenging their fundamental study or the testing of therapeutics. Notably, mouse bones do not reflect the human BM microenvironment thus precluding the possibility to investigate species-specific functional interactions between human cancer cells and their associated niches. The engineering of humanized ossicles (hOss) recently emerged as a promising technology capable of overcoming some of the mouse model limitations ^21–23^. These hOss are miniaturized bone organs offering the *in vivo* reconstitution of the human mesenchymal niche, providing a superior engraftment of human hematopoietic malignancies and some of the bone metastatic cancers ^21^. Beyond the heterogeneity of existing bioengineering protocols and their context-specific validation ^21^, an important limitation lies in the limited reproducibility of hOss approaches. In fact, over 90% of MSC donors fail to form hOss^19^. This considerable donor-to-donor variability combined with the limited MSCs ex vivo proliferation largely prevents the broad adoption of the hOss model by the community.

Hence, we here target the development of a standardized hOss system, offering the broad engraftment of patient-derived samples for healthy hematopoiesis, leukemia and solid tumor modeling. Our strategy relies on the potential identification and exploitation of a human mesenchymal cell line with intrinsic bone forming capacity. We previously established a telomerase-immortalized mesenchymal cell line (Mesenchymal Sword of Damocles, MSOD) from human primary BM-MSCs that allowed to overcome limited cellular proliferation, yet failed at generating hOss ^24^. Through constitutive BMP-2 implementation, the resulting MSOD-B line was recently exploited to engineer cell-free extracellular matrix (ECM) exhibiting frank osteoinductive properties ^25^. We hypothesized that MSOD-B cells can be implemented in a developmentally-inspired tissue engineering protocol to reproducibly generate hOss by endochondral (EC) and intramembranous (IM) ossifications. We postulate that the resulting “human mini-bones” will allow reconstituting a mesenchymal niche, permissive for human healthy and pathological hematopoiesis engraftment as well as supporting solid tumor metastasis.

## Results

### MSOD-B cells can reproducibly form distinct hOss

During embryonic development, bones are formed through EC and IM ossification processes. EC bone formation is characterized by MSCs differentiating into chondrocytes that produce cartilage as a precursor template to bone formation. Instead, IM ossification occurs without the involvement of a cartilage intermediate ^26^. MSOD-B cells were recently exploited to engineer cell-free osteoinductive ECM ^25^, yet their potential to recapitulate EC and IM programs towards the formation of ectopic bone organs remains to be determined. To this end, cells were seeded on collagen scaffolding material and exposed to either chondrogenic (Cho) or osteogenic (Ost) differentiation cues (**Fig.1a**). Following 3 weeks of *in vitro* differentiation, retrieved tissues displayed distinct cellular and ECM composition. At the cellular level, MSOD-B cells from Cho tissues exhibited a marked increase in intracellular levels of Sox9, collagen type 2 (Col2) and collagen type X (ColX) (**Fig.1b-c**), hallmarks of the Cho pathway activation. Contrary, cells from Ost tissues only displayed an increase in Sox9 level, however not associated with nuclear translocation (**Fig.1d**), indicating negligible Sox9 transcriptional activity. In line with the specific cellular profile, Cho tissues consisted in a glycosaminoglycans (**Fig.S1a-b**), Col2 and ColX-rich ECM (**Fig.1e**), a characteristic of mature hypertrophic cartilage templates. No cartilage-like features could be detected in Ost tissues (**Fig.1e** and **Fig.S1a**), which consisted of a mineralized collagen structure (**Fig.S1b**) resembling the osteoid matrix preceding bone formation.

**Fig.1.**
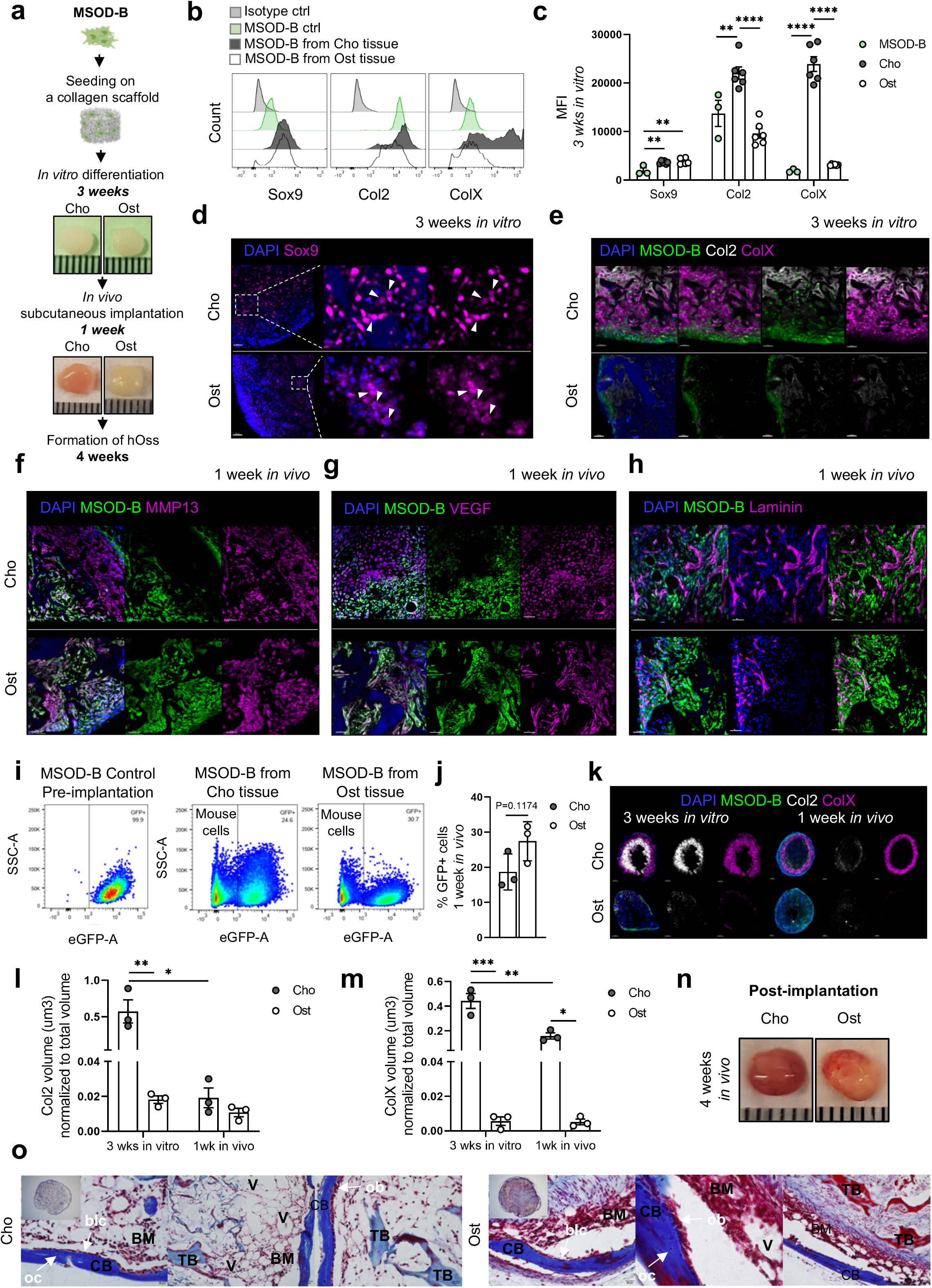
MSOD-B cells selectively primed towards EC and IM ossification pathways can reproducibly form hOss. **a,** Experimental scheme for *in vitro* and *in vivo* tissue generation. **b-c,** Representative flow cytometry histograms **(b)** and corresponding median fluorescence intensity (MFI) **(c)** of Sox9, Col2 and ColX staining performed on MSOD-B cells (pre-differentiation, control) and after isolation from Cho and Ost tissues 3-week post *in vitro* differentiation. Isotypes are also shown as control. Shown are mean values + 1 SE. n = 3-6 biological replicates. Statistical values were determined by one-way ANOVA followed by Tukey’s post-hoc test, **p < 0.01, ****p < 0.0001. **d,** Representative staining of Sox9 within Cho and Ost *in vitro* tissues. Nuclei are stained with DAPI. Arrowheads indicate Sox9 nuclear localization in MSOD-B cells within Cho tissues, while the cytoplasmic localization is shown for Ost tissues. n = 3 biological replicates. Scale bar = 50μm. **e,** Representative staining of Col2 and ColX within Cho and Ost *in vitro* tissues. Nuclei are stained with DAPI. n = 3 biological replicates. Scale bar = 50μm. **f-h,** Representative staining of MSOD-B, MMP13 **(f)**, VEGF **(g)** and Laminin **(h)** of Cho and Ost *in vivo* tissues explanted after 1-week of implantation. Nuclei are stained with DAPI. n = 3 biological replicates. Scale bar = 50μm. **i,** Representative flow cytometry plots of MSOD-B cells pre-implantation and isolated from Cho and Ost *in vivo* tissues 1-week post-implantation. **j,** Frequency (%) of MSOD-B cells within Cho and Ost tissues 1-week post-implantation. n = 3 biological replicates. Statistical values were determined by *t* test. **k,** Representative staining of MSOD-B, Col2 and ColX within 100μm thick Cho and Ost *in vitro* and *in vivo* tissue sections. Nuclei are stained with DAPI. n = 3 biological replicates. Scale bar = 500μm. **l-m**, Volume measurements of Col2 **(l)** and ColX **(m)** in μm^3^ estimated using Imaris software. Nuclei are stained with DAPI. n = 3 biological replicates. Statistical values were determined by two-way ANOVA followed by Šídák’s post-hoc test, *p < 0.05, **p < 0.01, ***p < 0.001. **n,** Representative macroscopic images of Cho and Ost hOss isolated at 4-week post-implantation. Size ≈ 0.5cm. **o**, Representative histological images of Masson’s trichrome staining of the sections from 4-week implanted Cho and Ost hOss. n = 3 biological replicates. BM (bone marrow), CB (cortical bone), TB (trabecular bone), V (vessels), blc (bone-lining cells), oc (osteocytes), ob (osteoblasts) shown with arrows.

Following *in vitro* engineering, Cho and Ost tissues were subcutaneously implanted in NOD.Cg-*Prkdc^scid^ Il2rg^tm1Wjl^*/SzJ (NSG) mice for further development (**Fig.1a**). As early as 1 week post-*in vivo* implantation, active remodeling of Cho and Ost tissues was evidenced by matrix metallopeptidase 13 (MMP13) and vascular endothelial growth factor (VEGF) staining (**Fig.1f-g** and **Fig.S1c-d**). This was associated with a rapid graft’s vascularization-in particular in Cho tissues-(**Fig.1h** and **Fig.S1e**) driven by invading mouse endothelial cells (**Fig.1i-j** and **Fig.S1f**). Using 3D quantitative microscopy imaging ^27^, we further report the presence but progressive cartilage digestion in Cho tissues *in vivo* (**Fig.1k-m** and **Fig.S1g**). In turn, cartilage was consistently absent in Ost tissues which accumulated osteocalcin (OCN) (**Fig.S1h-i**). Strikingly, at 4-week post-*in vivo* implantation both Cho and Ost grafts led to the successful formation of hOss (**Fig.1n**) with mature cortical and trabecular structures as well as marrow cavities (**Fig.1o**). Collectively, these data indicate that MSOD-B cells can successfully undergo Cho and Ost differentiation *in vitro*, forming mature hypertrophic cartilage and osteoid tissue respectively. Their subsequent ectopic implantation results in rapid hOss formation through recapitulation of distinct EC and IM bone developmental programs.

### MSOD-B hOss support the establishment of human hematopoiesis

In order to investigate whether MSOD-B hOss can support human hematopoiesis, NSG mice were implanted with Cho and Ost tissues and irradiated 4 weeks later for the intravenous (IV) transplantation of human cord blood CD34^+^ hematopoietic stem/progenitor cells (hCB-CD34+ HSPCs) (**Fig.2a**). The MSOD-B Cho and Ost implants development was first monitored over time by micro-computed tomography (μCT) at 4, 12 and 24-week post-implantation. This revealed a highly reproducible hOss formation with 96.25% of Cho and 96.7% of Ost tissues successfully developing into mature bone organs (>95 hOss assessed, **Fig.2b-d**). Important structural differences were observed between hOss types. Cho tissues developed into ~ 40mm^3^ organs (**Fig.S2a**) with a progressive decrease in bone volume (BV) (**Fig.2e** and **Fig.S2b**) resulting in a large BM cavity of high cellularity (**Fig.2f**). In contrast, Ost hOss appeared to be smaller organs (~20mm^3^) (**Fig.S2a**), predominantly composed of bone tissue with reduced BM space of significantly lower cellularity (**Fig.2e-f** and **Fig.S2b**).

**Fig.2.**
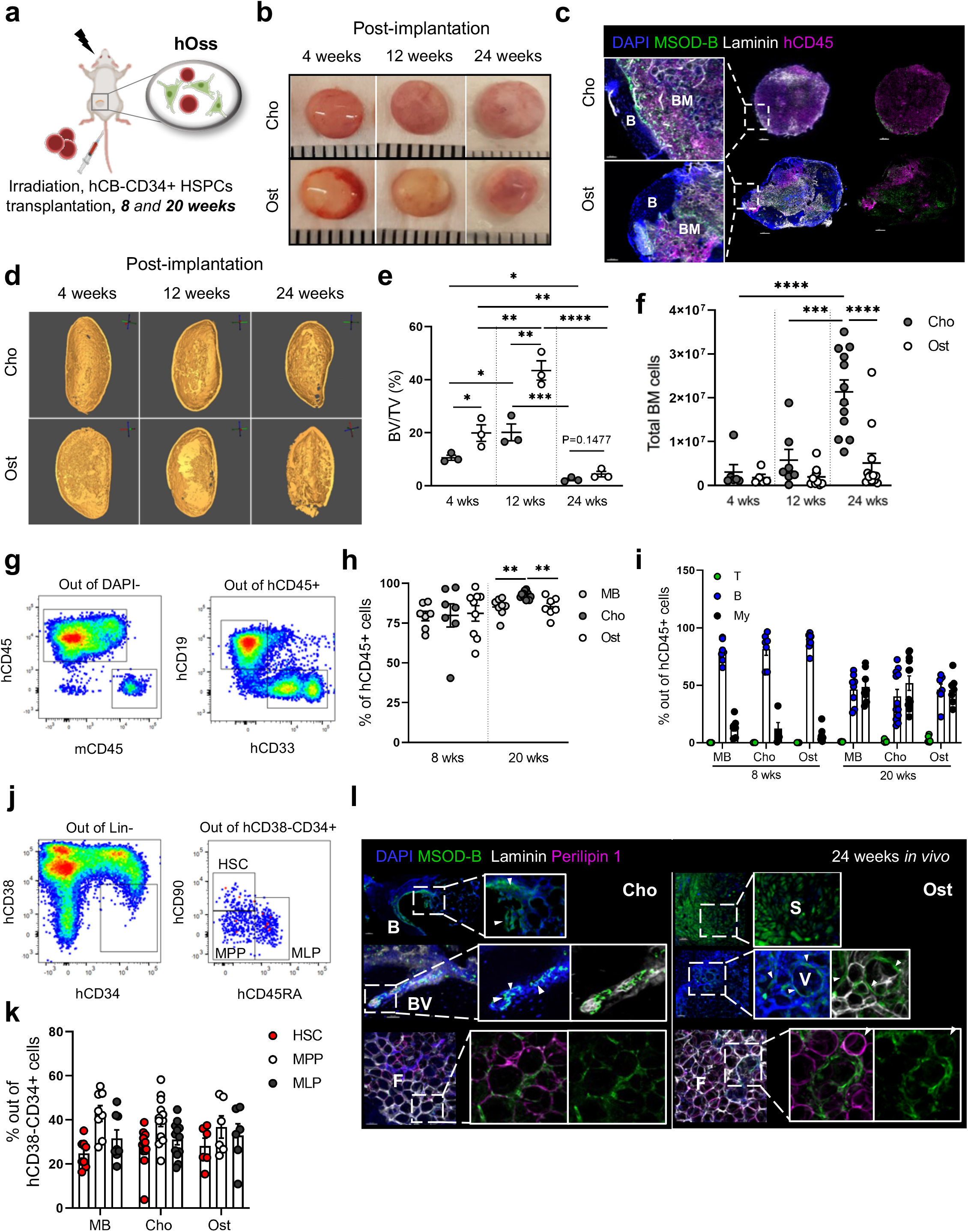
MSOD-B hOss reconstitute a human mesenchymal niche supporting the establishment of human hematopoiesis. **a,** Experimental scheme of CB-CD34^+^ cell transplantation. **b,** Representative macroscopic images of Cho and Ost hOss isolated at 4, 12 and 24-week post-implantation. Size ≈ 0.5cm. **c,** Representative staining of MSOD-B, hCD45^+^ cells and Laminin within Cho and Ost hOss explanted at 24-week post-implantation. Nuclei are stained with DAPI. n = 3 biological replicates. Scale bar = 500μm. **d,** μCT images of bones from Cho and Ost hOss explanted at 4, 12 and 24-week post-implantation. **e,** Quantification of bone volume (BV) normalized to total volume (TV) of Cho and Ost hOss explanted at 4, 12 and 24-week post-implantation. n = 3 biological replicates. Statistical values were determined by one-way ANOVA followed by Tukey’s post-hoc test, *p < 0.05, **p < 0.01, ***p < 0.001, ****p < 0.0001. **f,** Number of total BM cells isolated from Cho and Ost hOss at 4, 12 and 24-week post-implantation. n = 5-12 biological replicates. Statistical values were determined by one-way ANOVA followed by Tukey’s post-hoc test, **p < 0.01, ***p < 0.001. **g,** Schematic representation of the gating strategy of hCD45^+^ and mouse CD45^+^ (mCD45^+^), human CD19^+^ B cells and CD33^+^ myeloid (My) cells in Cho and Ost hOss explanted at 20-week post-transplantation. **h,** Percentage of hCD45+ cells in mouse bones (MB), Cho and Ost hOss at 8 and 20-week post-transplantation. n = 5–12 biological replicates. Statistical values were determined by one-way ANOVA followed by Tukey post-hoc test, **p < 0.01. **h-i,** Percentage of of hCD45^+^ **(h)**, CD3^+^ T, CD19^+^ B and CD33^+^ My cells out of hCD45^+^ cells **(i)** in MB, Cho and Ost hOss at 8 and 20-week post-transplantation. n = 7–12 biological replicates. Statistical values were determined by one-way ANOVA followed by Tukey’s post-hoc test, **p < 0.01. **j-k,** Schematic representation of the gating strategy **(j)** and percentage **(k)** of human CD38^-^CD34^+^, HSCs (CD38^-^CD34^+^CD90^+^CD45RA^-^), MLPs (CD38^-^CD34^+^CD90^-^CD45RA^+^) and MPPs (CD38^-^CD34^+^CD90^-^CD45RA^-^) in Cho and Ost hOss explanted at 20-week post-transplantation. n = 6–12 biological replicates. **l,** Representative staining of MSOD-B, Perilipin 1+ adipocytes and Laminin in Cho and Ost hOss at 24-week post-implantation. Nuclei are stained with DAPI. Scale bar = 50μm. n = 3 biological replicates. B (bone), BV (bone vessel), V (vessel), F (fat), S (stroma).

Flow cytometry analysis of mouse peripheral blood (PB) revealed a substantial engraftment of hCB-CD34^+^ cells, significantly increasing over time (**Fig.S2d-e**). Confocal analysis of explanted tissues indicated successful engraftment within both hOss types, forming human CD45^+^ (hCD45) hematopoietic islets (**Fig 2c** and **Fig.S2c)**. Quantitative analysis confirmed an overall very high engraftment level in Cho and Ost hOss (**Fig.2g-h**), as well as in mouse bone marrow (MB) (**Fig.2h**), with balanced lymphoid and myeloid lineage differentiation at 20-week post-transplantation (**Fig.2g, Fig.2i** and **Fig.S2e**). Noteworthy, Cho hOss showed a significantly higher engraftment compared to both Ost hOss and MB (**Fig.2h**). Primitive HSCs, multipotent progenitors (MPP) and multipotent lymphoid progenitors (MLP) were successfully identified in hOss and MB without marked frequency differences (**Fig.2j-k**).

Importantly, MSOD-B cells persisted in hOss and reconstituted a human mesenchymal niche in the form of bone-lining cells, adipocytes, stromal and perivascular cells (**Fig.2l** and **Fig.S2f**). Those MSOD-B cells could be retrieved from mature hOss following tissue digestion for further analysis (**Fig.S2g-i**). Cells displayed *in vitro* plastic adherence and proliferation capacity, while expressing the surface markers CD73, CD90 and CD105 (**Fig.S2j-l**). Remarkably, retrieved MSOD-B cells were able to form secondary hOss after 3 weeks of *in vitro* Cho differentiation and 4 weeks of *in vivo* implantation (**Fig.S2m-n**). This suggests the presence of MSOD-B cells with mesenchymal stem cell-like properties in engineered hOss ^28, 29^.

In summary, following EC and IM ossification routes MSOD-B cells can form distinct bone organs reconstituting a humanized mesenchymal niche, which supports the engraftment and differentiation of human CB-CD34^+^ cells. EC ossification resulted in a bigger bone marrow compartment of higher cellularity and human CB-CD34^+^ engraftment compared to IM ossification.

### AML cells preferentially home and engraft in MSOD-B hOss

A large number of primary human acute myeloid leukemia (AML) samples hardly engraft in patient-derived xenograft (PDX) mouse models. This prompted the development of advanced *in vivo* systems, including hOss ^30–34^ based on the human niche capacity to support leukemia survival and growth ^21–23^. Validating our MSOD-B hOss in this disease setting would thus provide an attractive solution towards the standardized modeling of AML *in vivo*.

Based on their higher BM cellularity and hematopoietic engraftment, Cho hOss were selected for assessing the engraftment of leukemic materials. Ten different patient’s BM aspirates were investigated, reflecting three clinical risk categories: favorable, intermediate and adverse ^35^ (**Fig.S3a**). NSG mice bearing hOss (or not, as control) were irradiated and transplanted with sorted CD45^+^CD19^-^CD3^-^CD34^+^ or CD45^+^CD19^-^CD3^-^ cells from AML primary samples (**Fig.3a**).

**Fig.3.**
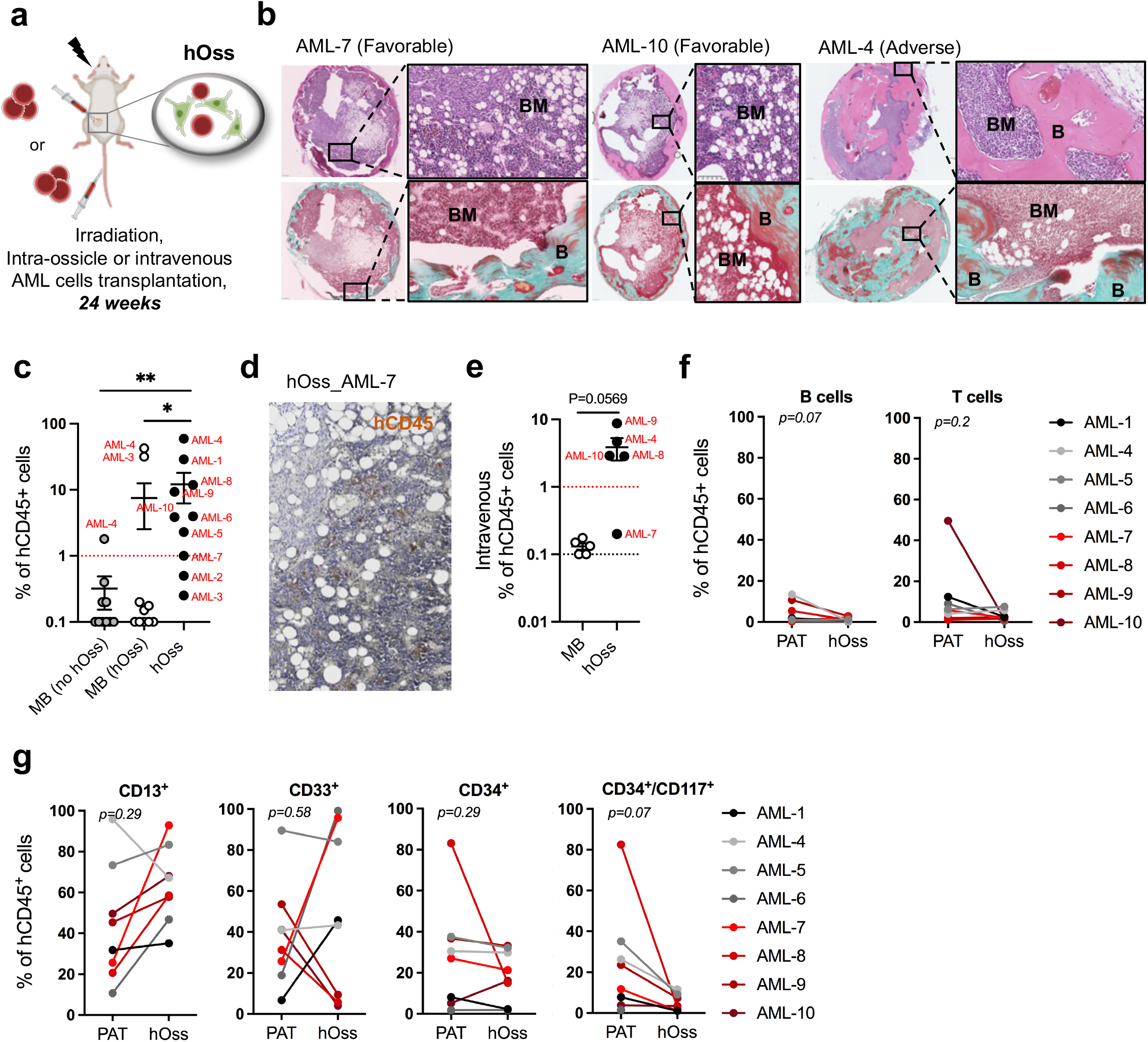
Primary acute myeloid leukemia cells preferentially home and engraft in MSOD-B hOss. **a,** Experimental scheme of AML patient sample transplantation. **b,** Representative histological images of Mayer’s Haematoxylin and Eosin (H&E) and Masson’s trichrome staining of the sections from AML-engrafted hOss. BM (bone marrow), B (bone). Scale bar = 250μm (whole section) and 50μm (zoomed areas). **c,** Percentage of hCD45^+^ leukemic cells in mouse bones (MB) of NSG mice not bearing hOss (MB no hOss, intravenous injection), in MB (hOss) and hOss at 24 weeks post-intra-ossicle transplantation. n = 11 NSG mice for intravenous transplantation; n = 2-4 biological replicates per AML patient sample for intra-ossicle transplantation, 10 AML patient samples in total. Statistical values were determined by Friedman’s test followed by Dunn’s post-hoc test, *p < 0.05, **p < 0.01. **d,** Representative staining of hCD45^+^ (terracotta) AML cells within hOss at 24-week post intra-ossicle transplantation. Scale bar = 50μm. **e,** Percentage of hCD45^+^ AML cells in MB and hOss at 24-week post-intravenous transplantation. n = 1-4 biological replicates per AML sample, 5 AML samples in total. Statistical values were determined by *t* test. **f-g,** Percentage of CD19^+^ B and CD3^+^ T cells (**f**), CD13^+^, CD33^+^ myeloid cells and CD34^+^, CD34^+^CD117^+^ (**g**) cells out of human CD45^+^ AML cells in patient (PAT) initial samples and hOss at 24-week post-intra-ossicle transplantation. n = 2–4 biological replicates per AML sample, 8 AML-engrafted samples in total. Statistical values were determined by paired *t* test.

All transplanted hOss successfully developed into mature bone organs with well-defined bone and bone marrow compartments (**Fig.3b**). The engraftment of AML samples was assessed at 24-week post-transplantation, and considered successful when the level of human CD45^+^ cells was higher than 1% of total BM mononuclear cells. AML engraftment was first assessed in mice not bearing hOss (MB-no hOss, **Fig.3c**), which resulted in only a single positive patient engraftment (AML-4). In sharp contrast, the transplantation of AML in hOss (intra-ossicle injection) led to 80% of sample engraftment in hOss with significant hCD45+ levels detected (**Fig.3c-d** and **Fig.S3c**). Limited engraftment was observed in PB, the spleen and MB (MB-hOss) of the very same mice (**Fig.3c** and **Fig.S3b**) with only 20% of AML samples passing the defined engraftment threshold. Intravenous transplantation was also performed using 5 AML samples, in order to assess a preferential homing to hOss over MB. Remarkably, while leukemic cells could not engraft in MB, most samples successfully engrafted in hOss with hCD45^+^ levels consistently superior to the ones detected in MB (**Fig.3e**).

A more detailed phenotypic analysis of engrafted AML patient cells in hOss was performed and compared to the initial patient (PAT) population frequencies. AML cells within hOss and the initial PAT samples displayed similar patterns with a poor lymphoid (CD19^+^ B and CD3^+^ T cells) (**Fig.3f** and **Fig.S3d**) and high myeloid reconstitution, being positive for CD13, CD33 as well as for CD34 and stem-like CD34/CD117 markers, characteristic of leukemic blasts ^36–38^ (**Fig.3g** and **Fig.S3d**). Together, we here demonstrate that the MSOD-B hOss microenvironment is permissive to the development of AML, which preferentially home and engraft in hOss when compared to mouse bone marrow.

### MSOD-B hOss support BC metastasis and NB cell engraftment

Bone is a privileged metastatic site for a large number of solid tumors ^17, 18, 39^. Deciphering mechanisms driving cancer metastasis would largely benefit from robust models capable of recapitulating tumor colonization to the BM microenvironment in a human specific fashion. To explore whether MSOD-B hOss can be exploited as a tumor model for bone colonization, we first assessed its capacity to act as a bone metastatic site in a breast cancer (BC) context. The aggressive MDA-MB-231 and the comparatively more lethargic MCF7 luciferase-labeled BC cell lines^40^ were intravenously injected into NSG mice bearing Cho hOss (**Fig.4a**). As previously reported ^41^, mice intravenously injected with MDA-MB-231 cells rapidly developed breathing problems caused by proliferating cancer cells in lungs, and therefore require euthanasia at 14-day post-injection. Despite the short *in vivo* period, 37.5% of hOss exhibited clear bioluminescence signal (**Fig.S4a-c**). Similarly, intravenously injected MCF7 cells also colonized the mouse lungs, and could subsequently be identified in hOss and mouse limbs at 21-day post-injection (**Fig.4b**). The combined *in vivo* bioluminescence and immunofluorescence analysis revealed that 62.5% of hOss were positively engrafted with MCF7 cells at 35-day post-transplantation (**Fig.4b-d**). Those data indicate that hOss can act as bone metastatic sites for human BC cells.

**Fig.4.**
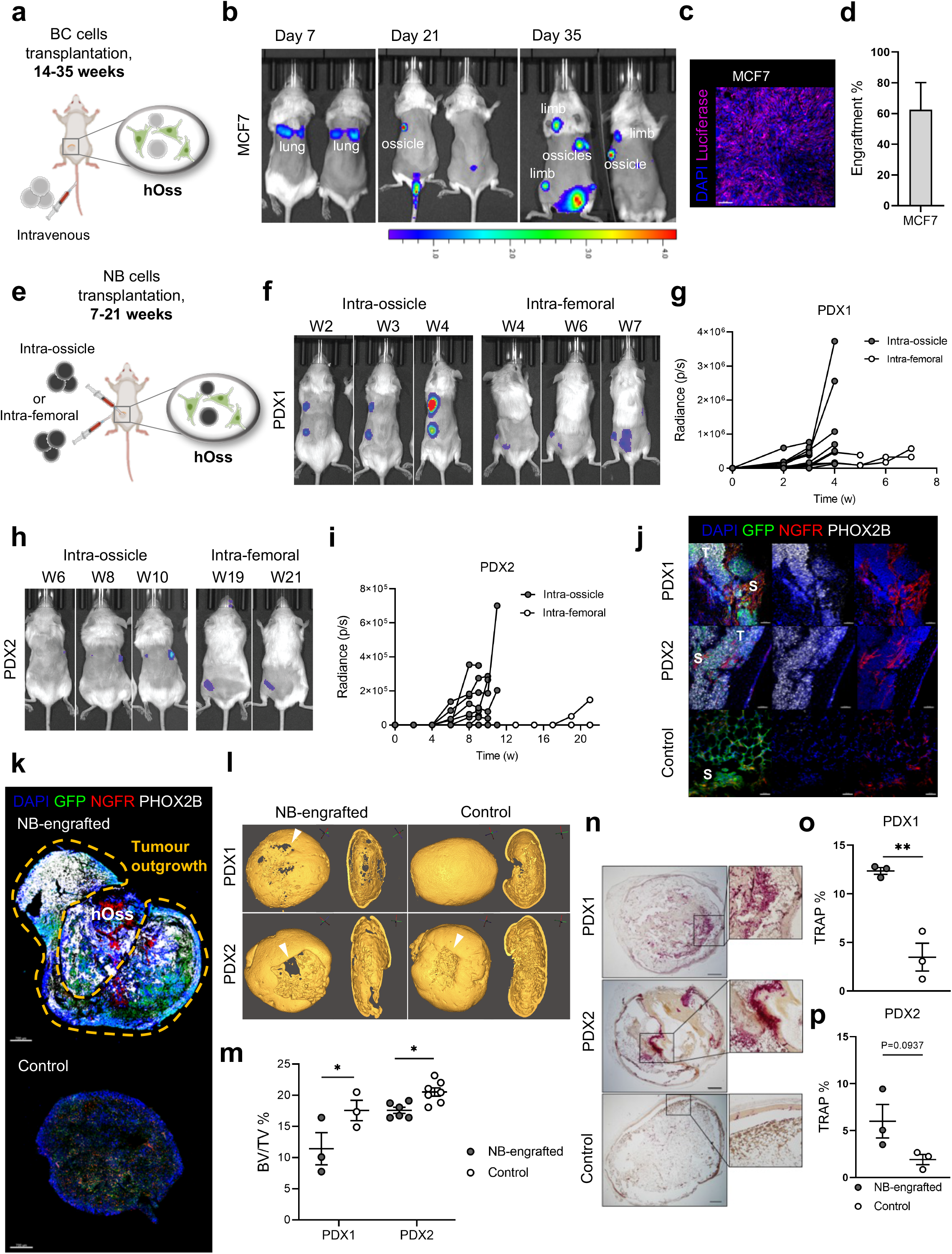
MSOD-B hOss support breast cancer metastasis and neuroblastoma cell engraftment. **a,** Experimental scheme of breast cancer (BC) cell transplantation. **b,** Representative bioluminescence images showing the *in vivo* distribution of MCF7 cells at 7, 21 and 35-day post-transplantation. **c,** Representative staining of Luciferase-MCF7 cells within hOss at 35-day post-transplantation. Nuclei are stained with DAPI. Scale bar = 50μm. **d,** Percentage of engrafted-hOss with MCF7 cells. n = 8 biological replicates. **e,** Schematic representation of the experimental setup of NB cell transplantation. **f,** Representative bioluminescence images showing the *in vivo* distribution of NB PDX1 cells within hOss and mouse femurs at 2-4-week (PDX1) and 4-7-week (femur) post-transplantation. **g,** Bioluminescence quantification (average radiance: photons/sec/cm^2^ in defined regions of interest) to monitor neuroblastoma (NB) PDX1 tumor burden in hOss and mouse femurs. n = 10 biological replicates for hOss, n = 3 mice for femurs. **h,** Representative bioluminescence images showing the *in vivo* distribution of NB PDX2 cells within hOss and mouse femurs at 6-10-week (PDX2) and 19-21-week (femur) post-transplantation. **i,** Bioluminescence quantification to monitor NB PDX2 tumor burden in hOss and mouse femurs. n = 10 biological replicates for hOss, n = 3 mice for femurs. **j-k,** Representative staining of GFP^+^ cells (MSOD-B and PDXs), NGFR (MSOD-B), PHOX2B (NB PDX1 and PDX2) in NB-engrafted and Control hOss. Nuclei are stained with DAPI. Yellow dashed line shows areas of tumor outgrowth in NB-engrafted hOss **(k)**. Scale bar = 50μm **(j)** and 700μm **(k)**. S (stroma), T (tumor). **l,** μCT images of bones from NB PDX1 and PDX2-engrafted and control hOss. Arrowheads indicate osteolytic lesions. **m,** Quantification of bone volume (BV) normalized to total volume (TV) of PDX1 and PDX2-engrafted and control hOss. n = 3-7 biological replicates. Statistical values were determined by multiple paired *t* test, *p < 0.05. **n-p,** Representative images **(n)** and quantification **(o-p)** of Tartrate-Resistant Acid Phosphatase (TRAP) staining for the detection of osteoclast activity in NB PDX1 **(o)** and NB PDX2 **(p)** engrafted and control hOss. n = 3 biological replicates. Scale bar = 500μm. Statistical values were determined by *t* test, **p < 0.01.

We next investigated the possibility to exploit MSOD-B hOss for neuroblastoma (NB), the most frequent extra-cranial pediatric malignancy, accounting for ~15% of all cancer-related deaths in children. NB primarily develops in the adrenal gland and rapidly metastasizes to other organs. Bone and marrow are the preferential metastatic sites, and the main reason for NB treatment relapse and mortality ^42^. While the NB cells-bone marrow niche interactions are postulated to be essential towards improving therapy success ^39, 43, 44^, currently no “human metastatic niche” models exist for *in vivo* studies in NB. To explore the MSOD-B hOss as a model for NB, two patient-derived xenograft (PDX) models, both derived from high-risk NB patients, were exploited: a primary tumor-derived (PDX1) and one derived from a brain metastasis (PDX2) ^45–47^. Intra-ossicle and intra-femoral injections of luciferase-labeled NB PDX1 and PDX2 cells were performed to evaluate NB cells engraftment and growth (**Fig.4e**). Bioluminescence signals validated the positive engraftment of PDXs cells following injection in mouse femurs and hOss (**Fig.4f-i** and **Fig.S4d**), which consistently remained confined and did not metastasize to peripheral organs or non-injected hOss. NB PDX1 cells engrafted in 90% of hOss and 100% of mice femurs, with a respective average engraftment time of 2-3 weeks versus 4-5 weeks (**Fig.4f-g**). NB PDX2 cells engrafted in 80% of hOss but in only 33% of femurs (**Fig.4h-i**). In line with PDX1 results, we observed a significantly faster PDX2 growth in hOss (9-weeks) versus femurs (19-weeks). Expression of NB markers paired-like homeobox 2b (PHOX2B) and neural cell adhesion molecule CD56 confirmed the presence of NB cell in hOss, while nerve growth factor receptor^+^ (NGFR)/PHOX2B^-^ phenotype evidenced MSOD-B cells (**Fig.4j-k** and **Fig.S4e**). Flow cytometry analysis of NB cell frequencies indicated 41.1% and 28.86% of GD2^+^ fraction in PDX1 and PDX2-engrafted hOss respectively. In addition, a GD2^+^CD56^+^ fraction was also identified (48.83% for PDX1, 33.52% for PDX2) (**Fig.S4f-g**). Through 3D scanning of large tissue sections, a spectacular tumor development was evidenced in injected hOss, with presence of NB cells both within the bone marrow cavity as well as in the form of an outgrown mass engulfing the whole cortical structure (**Fig.4k** and **Fig.S4h**). Strikingly, μCT divulged the presence of bone lesions in PDX1 and PDX2-engrafted hOss, resulting in a quantifiable bone loss (**Fig.4l-m**). Since NB cells were previously reported capable of recruiting osteoclasts in bone matrix ^48^, we performed tartrate-resistant acid phosphatase (TRAP) staining of hOss (**Fig.4n-p**). This allowed identifying a marked increase in osteoclastic activity in NB-engrafted hOss, as compared to control ones. This data suggests the recapitulation of the osteolytic disease-associated pattern, leading to skeletal alterations in neuroblastoma patients.

## Discussion

We here report a versatile and reliable hOss model supporting the establishment of human hematopoiesis, the engraftment of primary AML samples, and acting as metastatic organ for solid tumor development.

Preexisting hOss approaches demonstrated the functional advantage of engineering a human BM niche ^21^. Thus far, however, protocols have remained relatively complex as well as requiring the use of primary MSCs, challenging their reproducibility. Instead, the MSOD-B consists in a mesenchymal cell line conserving lineage plasticity, capable of undergoing EC and IM ossifications. The process is highly efficient and in appearance limitless, with the same clonal population exploited for the engineering of over 1000 hOss. The 3 weeks *in vitro* priming followed by *in vivo* implantation led to the formation of bone organs as early as 4-week post-implantation, harboring a mature architecture comparable to native bones and composed of a functional human mesenchymal niche. In comparison, selected batches of primary MSCs previously required 5 weeks of *in vitro* differentiation to reach hypertrophy, and 8 to 12 weeks of *in vivo* development ^49^.

Beyond standardization, the MSOD-B line will also facilitate the generation of customized hOss carrying gain/loss-of-function of specific genes ^50, 51^. CRISPR/Cas9 may be used to interrogate molecular pathways potentially involved in the maintenance of specific hematopoietic or cancer populations. Conversely, additional transgene could be implemented and conditionally expressed, thus providing an *in vivo* platform for probing candidate niche factors. Their overexpression and presentation by human niche cells may even further increase engraftment, as a credible alternative to current mouse engineering techniques ^52^. In fact, while MSOD-B hOss showed clear superiority to NSG mice with broad primary AML engraftment, some patient samples did not engraft. Extensive *in vivo* time may be required for the engraftment of certain AML subtypes ^53^. It warrants further investigation, together with the capacity to not only maintain primary clonal architecture, but ideally also mimic their natural evolution patterns. This is however challenged by the difficult access to patients’ biopsies. Of importance, the successful engraftment reported in MSOD-B hOss was achieved using a lower amount of AML cells as compared to what is routinely used for generating AML PDXs ^54, 55^. This indicates that the hOss model may also reveal advantageous for patient materials of limited quantity.

Our findings prompt also investigating the suitability of MSOD-B hOss in other pathological contexts. This is of particular interest for malignancies hardly developing in mice, such as myelodysplastic syndromes which successful engraftment highly depends on the local microenvironment ^56^. Similarly, solid cancers are ideal candidates in light of the severe prognosis typically associated with bone and BM metastasis. In the present study, we demonstrate successful colonization of hOss by BC and NB tumor cells after intravenous and intra-ossicle injections, respectively. The BM was previously pointed out as metastatic site primarily responsible for treatment resistance and relapse of several cancers ^43, 44, 57^. The recapitulation of some of the disease-specific patterns such as osteolytic lesions in NB holds great promise, towards exploiting MSOD-B hOss as advanced pre-clinical system. Through a parallel monitoring of human cancer and niche cells, one may shade light on how cancer cells can reprogram their local environment to develop resistance to chemotherapy ^39, 57^ or enforce their dissemination ^58^. The subsequent testing of therapeutics in a personalized fashion, may help identifying substances to inhibit cancer metastasis.

Limitations of the MSOD-B hOss principally lie in its chimeric nature, including a vasculature of mouse origin. In addition, while cell lines confer standardization, their biological relevance may not fully match with their primary counterparts. This prompts additional study, including the possibility of further humanization of the model.

Collectively, the MSOD-B hOss consist in malleable human mini-bones to be used as a fundamental or pre-clinical *in vivo* platform. The standardization of such humanized systems may reveal essential towards compiling human-specific findings, and accelerate the design of personalized therapies.

## Author contributions

P.E.B. conceptualized the study. A.G. and P.E.B. designed the experiments and interpreted the results. A.G. contributed to all experiments described in the manuscript. A.B., C.RM.C., AK.F. and B.P. carried out the AML study. D.Z., K.A., A.M., J.E. and D.B. carried out the NB study. D.Z., P.K., E.E. and C.D.M. carried out the BC study. N.K. and S.P. contributed to histological staining. A.G.G. performed μCT analysis. A.G. wrote the initial draft. A.G. and P.E.B. wrote the final manuscript. All authors contributed to the manuscript editing.

## Acknowledgments

We thank Lund Stem Cell Center FACS Facility and Imaging Facility and the Lund University Bioimaging Center for X-ray CT system. Lund University and Medicon village are gratefully acknowledged for providing IVIS-CT experimental resources. BioRender.com was used to present the experimental setups. A.G., D.Z., A.G.G., S.P. and P.E.B. were supported by the Knut and Alice Wallenberg foundation, the Medical Faculty at Lund University and Region Skåne. We also thank the Crafoordska stiftelsen for financial support (to P.E.B). This project has received funding from the European Research Council (ERC) (Starting grant hOssicle #948588 to P.E.B). A.G. was supported by The Royal Physiographic Society of Lund. K.A., A.M., J.E. and D.B. were supported by the Swedish Cancer Society and the Swedish Childhood Cancer Foundation. Work in the Porse lab was supported by the Danish Research Center for Precision Medicine in Blood Cancers (Danish Cancer Society grant no. R223-A13071), the Greater Copenhagen Health Science Partners and through a centre grant from the Novo Nordisk Foundation (Novo Nordisk Foundation Centre for Stem Cell Biology, DanStem; Grant Number NNF17CC0027852). E.E was supported by the Swedish Society for Medical Research (EE), Royal Physiographic Society of Lund (2019, EE). C.D.M. and P.K. were supported by BioCARE, Sweden. C.D.M was supported by the Ragnar Söderberg Foundation (N91/15), Cancerfonden (CAN 2016/783, CAN 2018/230 and 19 0632 Pj), and Swedish Research Council (2017-03389, 2019-02355 and 2020-02088).

## References

1. Friedenstein, A.J., Chailakhyan, R.K. & Gerasimov, U.V. Bone marrow osteogenic stem cells: in vitro cultivation and transplantation in diffusion chambers. Cell Tissue Kinet 20, 263–272 (1987).

2. Pittenger, M.F. et al. Multilineage potential of adult human mesenchymal stem cells. Science (New York, N.Y.) 284, 143–147 (1999).

3. Salhotra, A., Shah, H.N., Levi, B. & Longaker, M.T. Mechanisms of bone development and repair. Nature Reviews Molecular Cell Biology (2020).

4. Frenette, P.S., Pinho, S., Lucas, D. & Scheiermann, C. Mesenchymal stem cell: keystone of the hematopoietic stem cell niche and a stepping-stone for regenerative medicine. Annual review of immunology 31, 285–316 (2013).

5. Eaves, C.J. Hematopoietic stem cells: concepts, definitions, and the new reality. Blood 125, 2605–2613 (2015).

6. Comazzetto, S., Shen, B. & Morrison, S.J. Niches that regulate stem cells and hematopoiesis in adult bone marrow. Developmental cell 56, 1848–1860 (2021).

7. Mendez-Ferrer, S. et al. Mesenchymal and haematopoietic stem cells form a unique bone marrow niche. Nature 466, 829–834 (2010).

8. Calvi, L.M. et al. Osteoblastic cells regulate the haematopoietic stem cell niche. Nature 425, 841–846 (2003).

9. Kumar, S. & Geiger, H. HSC Niche Biology and HSC Expansion Ex Vivo. Trends in molecular medicine 23, 799–819 (2017).

10. Woods, K. & Guezguez, B. Dynamic Changes of the Bone Marrow Niche: Mesenchymal Stromal Cells and Their Progeny During Aging and Leukemia. Frontiers in cell and developmental biology 9 (2021).

11. Kode, A. et al. Leukaemogenesis induced by an activating β-catenin mutation in osteoblasts. Nature 506, 240–244 (2014).

12. Raaijmakers, M.H. et al. Bone progenitor dysfunction induces myelodysplasia and secondary leukaemia. Nature 464, 852–857 (2010).

13. de Jong, M.M.E. et al. The multiple myeloma microenvironment is defined by an inflammatory stromal cell landscape. Nature immunology 22, 769–780 (2021).

14. Gundem, G. et al. The evolutionary history of lethal metastatic prostate cancer. Nature 520, 353–357 (2015).

15. Kennecke, H. et al. Metastatic behavior of breast cancer subtypes. J Clin Oncol 28, 3271–3277 (2010).

16. Smid, M. et al. Subtypes of breast cancer show preferential site of relapse. Cancer research 68, 3108–3114 (2008).

17. Hochheuser, C. et al. The Metastatic Bone Marrow Niche in Neuroblastoma: Altered Phenotype and Function of Mesenchymal Stromal Cells. Cancers 12 (2020).

18. Zhang, W. et al. The bone microenvironment invigorates metastatic seeds for further dissemination. Cell 184, 2471–2486.e2420 (2021).

19. Côme, C., Balhuizen, A., Bonnet, D. & Porse, B.T. Myelodysplastic syndrome patient-derived xenografts: from no options to many. Haematologica 105, 864–869 (2020).

20. Kuperwasser, C. et al. A mouse model of human breast cancer metastasis to human bone. Cancer research 65, 6130–6138 (2005).

21. Dupard, S.J., Grigoryan, A., Farhat, S., Coutu, D.L. & Bourgine, P.E. Development of Humanized Ossicles: Bridging the Hematopoietic Gap. Trends in molecular medicine 26, 552–569 (2020).

22. Abarrategi, A. et al. Modeling the human bone marrow niche in mice: From host bone marrow engraftment to bioengineering approaches. The Journal of experimental medicine 215, 729–743 (2018).

23. Pievani, A. et al. Harnessing Mesenchymal Stromal Cells for the Engineering of Human Hematopoietic Niches. Front Immunol 12, 631279 (2021).

24. Bourgine, P. et al. Combination of immortalization and inducible death strategies to generate a human mesenchymal stromal cell line with controlled survival. Stem cell research 12, 584–598 (2014).

25. Pigeot, S. et al. Manufacturing of Human Tissues as off-the-Shelf Grafts Programmed to Induce Regeneration. Adv Mater, e2103737 (2021).

26. Kronenberg, H.M. Developmental regulation of the growth plate. Nature 423, 332–336 (2003).

27. Coutu, D.L., Kokkaliaris, K.D., Kunz, L. & Schroeder, T. Multicolor quantitative confocal imaging cytometry. Nature methods 15, 39–46 (2018).

28. Sacchetti, B. et al. Self-renewing osteoprogenitors in bone marrow sinusoids can organize a hematopoietic microenvironment. Cell 131, 324–336 (2007).

29. Dominici, M. et al. Minimal criteria for defining multipotent mesenchymal stromal cells. The International Society for Cellular Therapy position statement. Cytotherapy 8, 315–317 (2006).

30. Vaiselbuh, S.R., Edelman, M., Lipton, J.M. & Liu, J.M. Ectopic human mesenchymal stem cell-coated scaffolds in NOD/SCID mice: an in vivo model of the leukemia niche. Tissue engineering. Part C, Methods 16, 1523–1531 (2010).

31. Reinisch, A. et al. A humanized bone marrow ossicle xenotransplantation model enables improved engraftment of healthy and leukemic human hematopoietic cells. Nature medicine 22, 812–821 (2016).

32. Abarrategi, A. et al. Versatile humanized niche model enables study of normal and malignant human hematopoiesis. The Journal of clinical investigation 127, 543–548 (2017).

33. Pievani, A. et al. Acute myeloid leukemia shapes the bone marrow stromal niche in vivo. Haematologica (2020).

34. Antonelli, A. et al. Establishing human leukemia xenograft mouse models by implanting human bone marrow-like scaffold-based niches. Blood 128, 2949–2959 (2016).

35. Herold, T. et al. Validation and refinement of the revised 2017 European LeukemiaNet genetic risk stratification of acute myeloid leukemia. Leukemia 34, 3161–3172 (2020).

36. Casasnovas, R.O. et al. Immunological classification of acute myeloblastic leukemias: relevance to patient outcome. Leukemia 17, 515–527 (2003).

37. Gorczyca, W. et al. Immunophenotypic pattern of myeloid populations by flow cytometry analysis. Methods Cell Biol 103, 221–266 (2011).

38. Wells, S.J., Bray, R.A., Stempora, L.L. & Farhi, D.C. CD117/CD34 expression in leukemic blasts. Am J Clin Pathol 106, 192–195 (1996).

39. Hochheuser, C. et al. Mesenchymal Stromal Cells in Neuroblastoma: Exploring Crosstalk and Therapeutic Implications. Stem cells and development 30, 59–78 (2021).

40. Wang, H. et al. The osteogenic niche promotes early-stage bone colonization of disseminated breast cancer cells. Cancer cell 27, 193–210 (2015).

41. Minn, A.J. et al. Genes that mediate breast cancer metastasis to lung. Nature 436, 518–524 (2005).

42. Tolbert, V.P. & Matthay, K.K. Neuroblastoma: clinical and biological approach to risk stratification and treatment. Cell and tissue research 372, 195–209 (2018).

43. Seeger, R.C. et al. Quantitative tumor cell content of bone marrow and blood as a predictor of outcome in stage IV neuroblastoma: a Children’s Cancer Group Study. J Clin Oncol 18, 4067–4076 (2000).

44. Meads, M.B., Hazlehurst, L.A. & Dalton, W.S. The bone marrow microenvironment as a tumor sanctuary and contributor to drug resistance. Clin Cancer Res 14, 2519–2526 (2008).

45. Braekeveldt, N. et al. Neuroblastoma patient-derived orthotopic xenografts retain metastatic patterns and geno- and phenotypes of patient tumours. International journal of cancer 136, E252–261 (2015).

46. Persson, C.U. et al. Neuroblastoma patient-derived xenograft cells cultured in stem-cell promoting medium retain tumorigenic and metastatic capacities but differentiate in serum. Scientific reports 7, 10274 (2017).

47. Braekeveldt, N. et al. Patient-Derived Xenograft Models Reveal Intratumor Heterogeneity and Temporal Stability in Neuroblastoma. Cancer research 78, 5958–5969 (2018).

48. Sohara, Y. et al. Lytic bone lesions in human neuroblastoma xenograft involve osteoclast recruitment and are inhibited by bisphosphonate. Cancer research 63, 3026–3031 (2003).

49. Scotti, C. et al. Engineering of a functional bone organ through endochondral ossification. Proceedings of the National Academy of Sciences of the United States of America 110, 3997–4002 (2013).

50. Bourgine, P.E. et al. Fate Distribution and Regulatory Role of Human Mesenchymal Stromal Cells in Engineered Hematopoietic Bone Organs. iScience 19, 504–513 (2019).

51. Carretta, M. et al. Genetically engineered mesenchymal stromal cells produce IL-3 and TPO to further improve human scaffold-based xenograft models. Experimental hematology 51, 36–46 (2017).

52. Yin, L., Wang, X.J., Chen, D.X., Liu, X.N. & Wang, X.J. Humanized mouse model: a review on preclinical applications for cancer immunotherapy. Am J Cancer Res 10, 4568–4584 (2020).

53. Paczulla, A.M. et al. Long-term observation reveals high-frequency engraftment of human acute myeloid leukemia in immunodeficient mice. Haematologica 102, 854–864 (2017).

54. Hassan, N., Yang, J. & Wang, J.Y. An Improved Protocol for Establishment of AML Patient-Derived Xenograft Models. STAR Protoc 1, 100156 (2020).

55. Sanchez, P.V. et al. A robust xenotransplantation model for acute myeloid leukemia. Leukemia 23, 2109–2117 (2009).

56. Mian, S.A. et al. Ectopic humanized mesenchymal niche in mice enables robust engraftment of myelodysplastic stem cells. Blood Cancer Discov 2, 135–145 (2021).

57. Boutin, L. et al. Mesenchymal stromal cells confer chemoresistance to myeloid leukemia blasts through Side Population functionality and ABC transporter activation. Haematologica 105, 987–9998 (2020).

58. Shiozawa, Y., Eber, M.R., Berry, J.E. & Taichman, R.S. Bone marrow as a metastatic niche for disseminated tumor cells from solid tumors. Bonekey Rep 4, 689 (2015).

