## Supplementary Figures for "Engineering of human mini-bones for the standardized modeling of healthy hematopoiesis, leukemia and solid tumor metastasis"

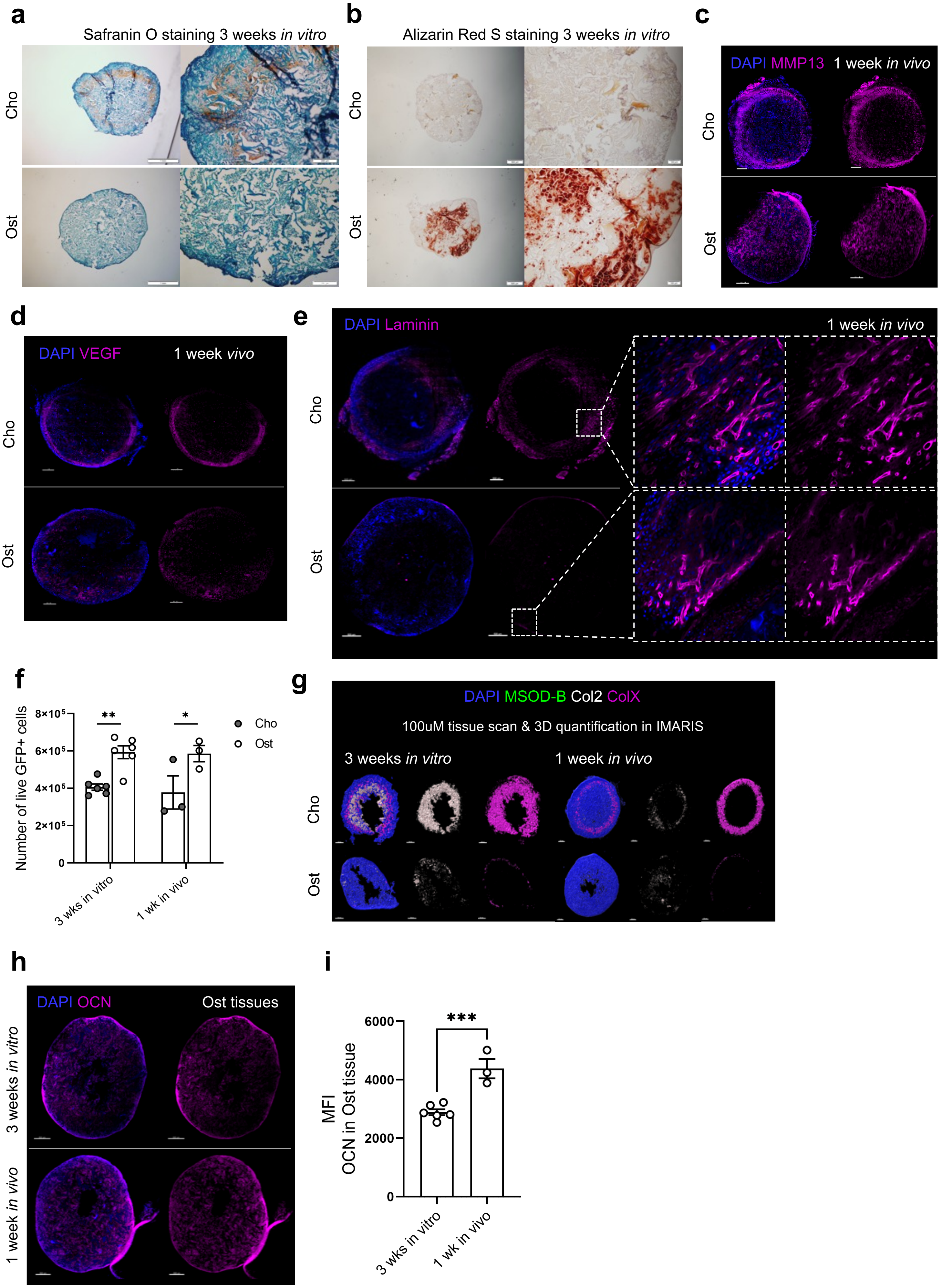

a
Safranin O staining 3 weeks in vitro
Cho
Ost
b
Alizarin Red S staining 3 weeks in vitro
Cho
Ost
c
DAPI MMP13 1 week in vivo
Cho
Ost
d
DAPI VEGF 1 week vivo
Cho
Ost
e
DAPI Laminin 1 week in vivo
Cho
Ost
f
Number of live GFP+ cells
8×10⁵
6×10⁵
4×10⁵
2×10⁵
0
**
*
Cho
Ost
3 wks in vitro
1 wk in vivo
g
DAPI MSOD-B Col2 ColX
100uM tissue scan & 3D quantification in IMARIS
3 weeks in vitro
1 week in vivo
Cho
Ost
h
DAPI OCN Ost tissues
3 weeks in vitro
1 week in vivo
i
MFI
OCN in Ost tissue
6000
4000
2000
0
***
3 wks in vitro
1 wk in vivo


Fig.S1

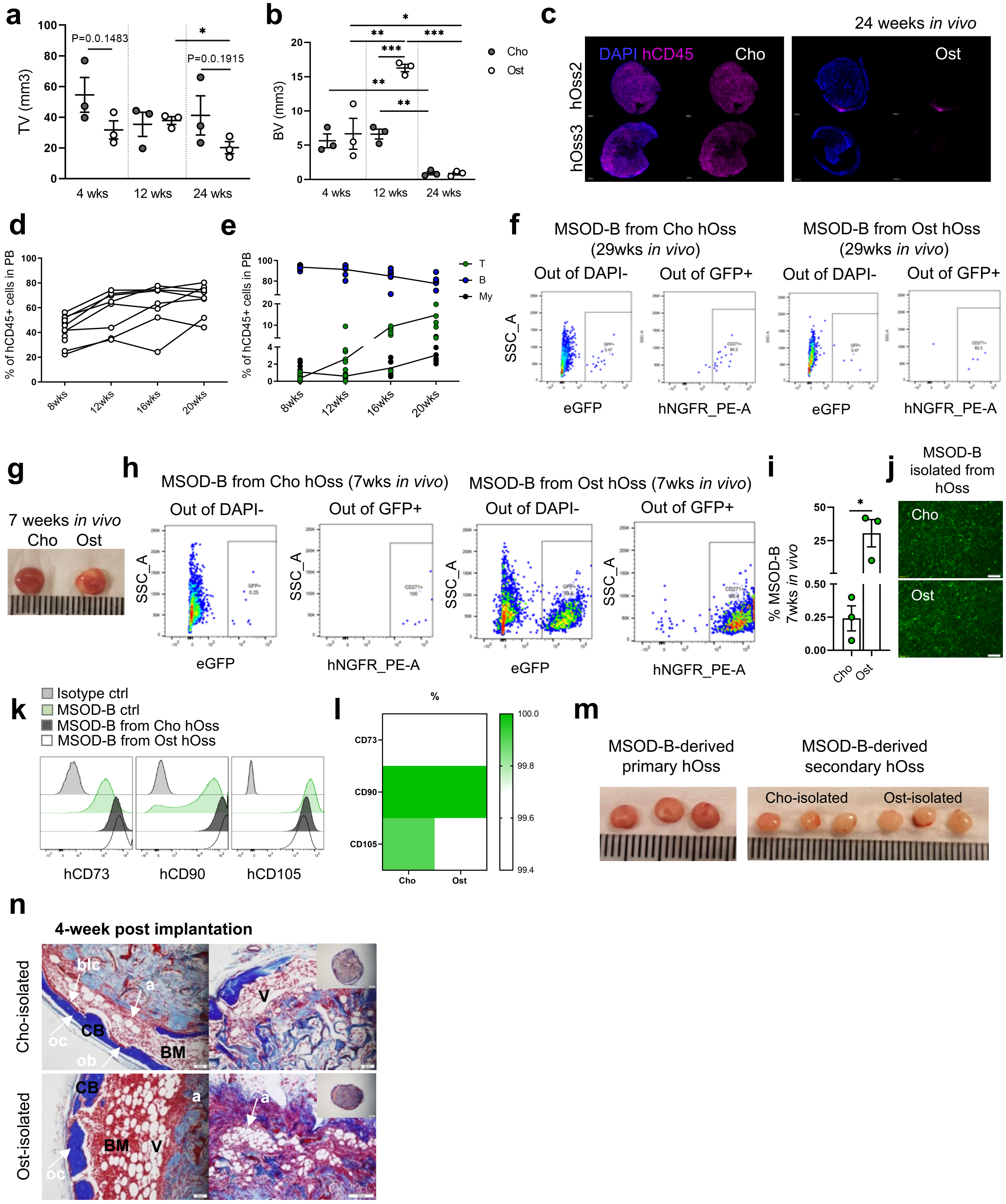

a
TV (mm3)
P=0.0.1483
P=0.0.1915
4 wks
12 wks
24 wks
b
BV (mm3)
Cho
Ost
c
24 weeks in vivo
DAPI hCD45
hOss2
hOss3
d
% of hCD45+ cells in PB
8wks
12wks
16wks
20wks
e
T
B
My
f
MSOD-B from Cho hOss (29wks in vivo)
MSOD-B from Ost hOss (29wks in vivo)
Out of DAPI-
Out of GFP+
SSC_A
eGFP
hNGFR_PE-A
g
7 weeks in vivo
h
MSOD-B from Cho hOss (7wks in vivo)
MSOD-B from Ost hOss (7wks in vivo)
i
% MSOD-B 7wks in vivo
j
MSOD-B isolated from hOss
k
Isotype ctrl
MSOD-B ctrl
MSOD-B from Cho hOss
MSOD-B from Ost hOss
hCD73
hCD90
hCD105
l
%
CD73
CD90
CD105
m
MSOD-B-derived primary hOss
MSOD-B-derived secondary hOss
Cho-isolated
Ost-isolated
n
4-week post implantation
Cho-isolated
Ost-isolated
blc
a
CB
oc
ob
BM
V


Fig.S2

**a**

| Sample | Age [y] | Disease stage | Diagnosis | FAB type | WHO classification | ELN risk group | Blasts in BM [%] | Gene | Gene | Gene | Gene | Sorted cells |
|---|---|---|---|---|---|---|---|---|---|---|---|---|
| AML-1 | 73 | Initial | AML | M4 | not otherwise categorized | Intermediate | 51 | DNMT3A | NPM1 | | | CD45+/CD3-/CD19- |
| AML-2 | 53 | Initial | AML | M0 | recurrent genetic abnormalities, i.e.with specific chromosomal changes | Adverse | 74 | CEBPA | NRAS | RUNX1 | ETV6 | CD45+/CD3-/CD19-/CD34+ |
| AML-3 | NA | Initial | AML | M4 | recurrent genetic abnormalities, i.e. with specific chromosomal changes | Favorable | 56 | CEBPA | NRAS | GATA2 | | CD45+/CD3-/CD19-/CD34+ |
| AML-4 | NA | Initial | AML | M2 | myelodysplasia-related changes, or abnormalities in how the blood cells look | Adverse | 31 | KRAS | NRAS | FLT3 | | CD45+/CD3-/CD19-/CD34+ |
| AML-5 | 32 | Initial | AML | M1 | not otherwise categorized | Adverse | 75 | NF1 | | | | CD45+/CD3-/CD19-/CD34+ |
| AML-6 | 56 | Initial | AML | M5 | not otherwise categorized | Favorable | 55 | NPM1 | | | | CD45+/CD3-/CD19- |
| AML-7 | 73 | Initial | AML | M4 | not otherwise categorized | Adverse | 83 | NA | | | | CD45+/CD3-/CD19- |
| AML-8 | 65 | Initial | AML | M0 | not otherwise categorized | Intermediate | 66 | DNMT3A | IDH2 | | | CD45+/CD3-/CD19- |
| AML-9 | 68 | Initial | AML | M2 | not otherwise categorized | Intermediate | 31 | SF3B1 | | | | CD45+/CD3-/CD19- |
| AML-10 | 56 | Initial | AML | M0 | recurrent genetic abnormalities, i.e. with specific chromosomal changes | Favorable | 85 | ETV6 | NPM1 | WT1 | PHF6 | CD45+/CD3-/CD19- |

French-American-British classification system (FAB); EuropeanLeukemiaNet classification system (ELN); Bone marrow (BM); Not available (NA)

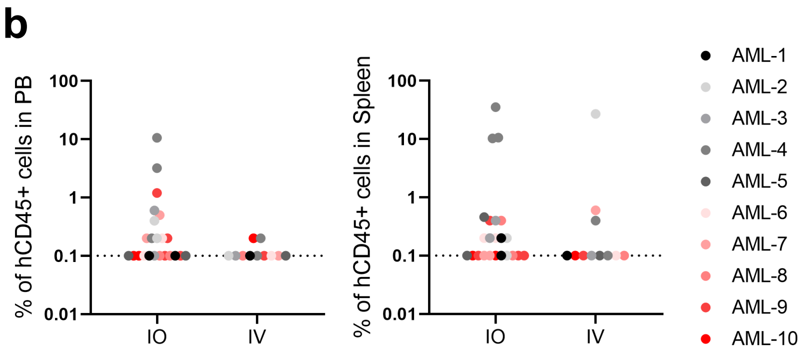


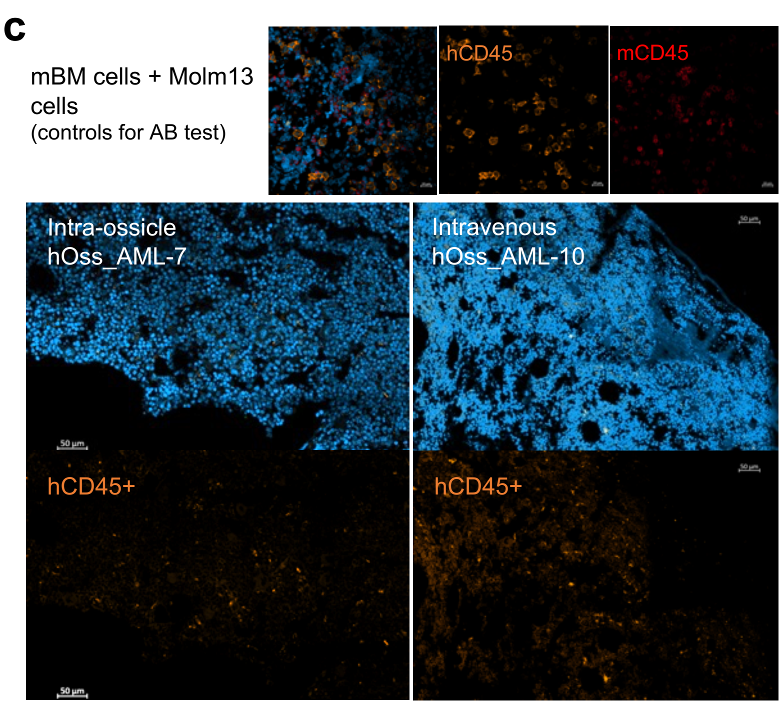


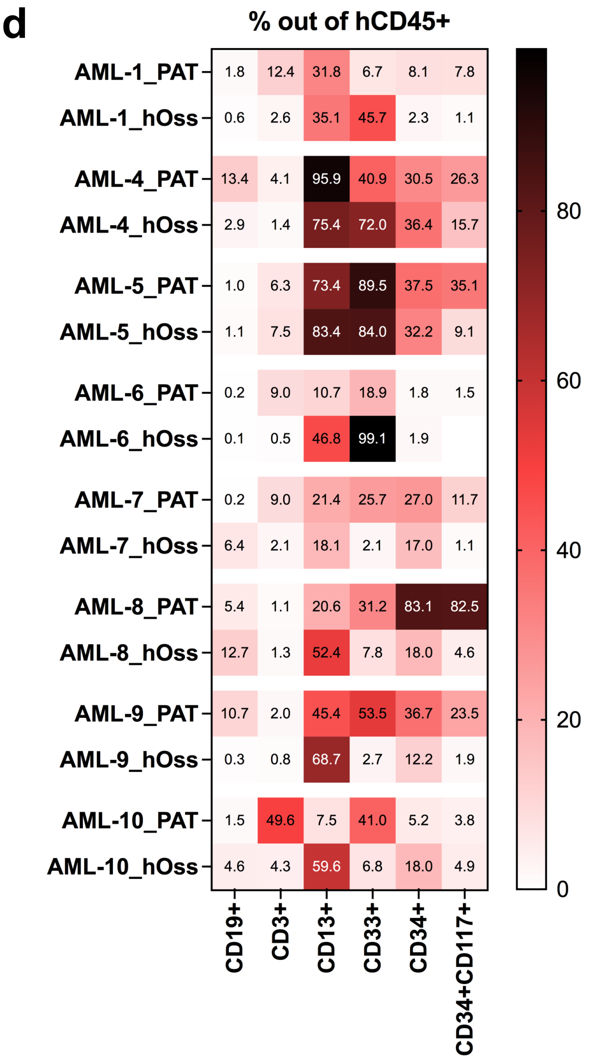


Fig.S3

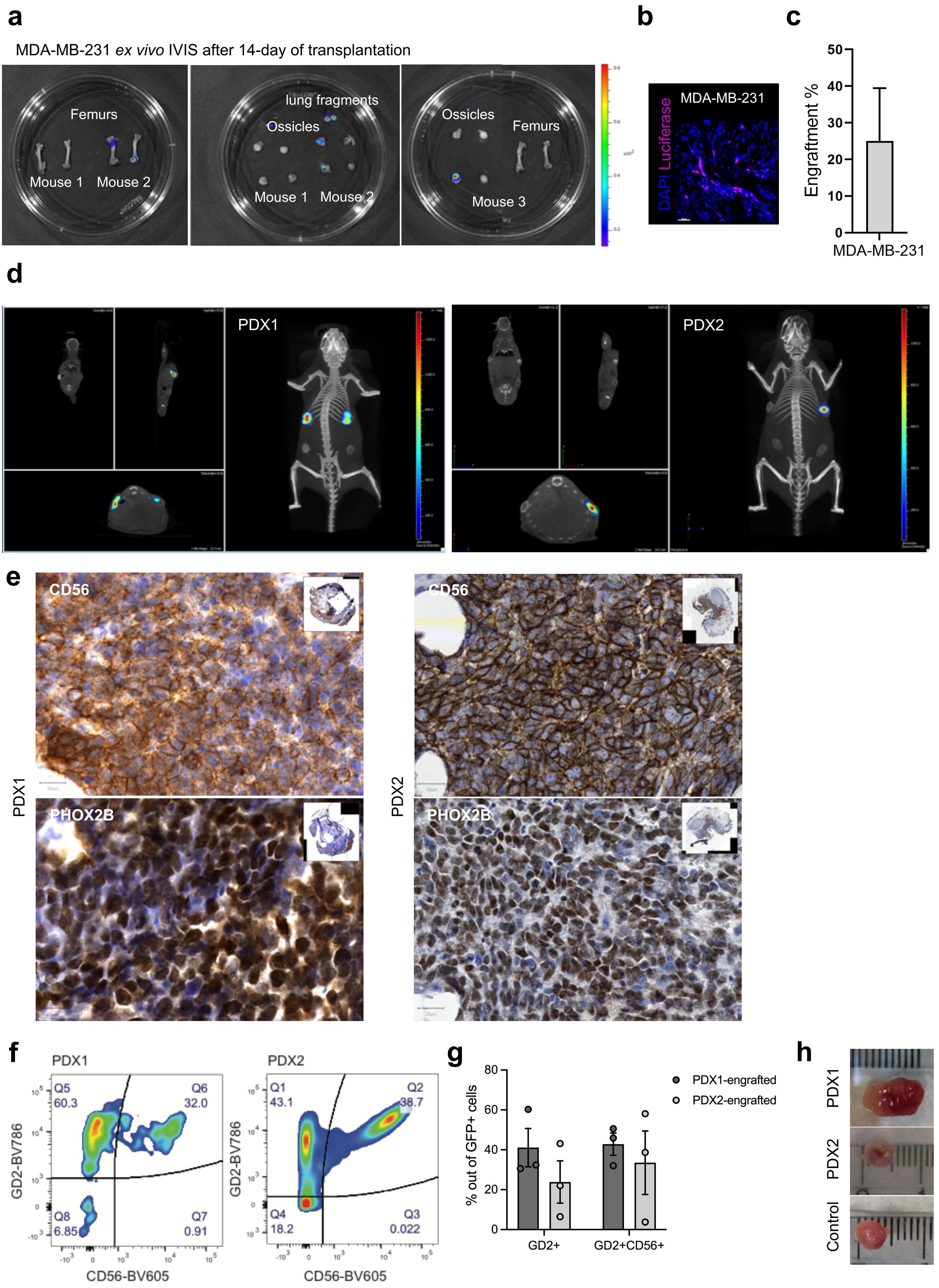
a
MDA-MB-231 ex vivo IVIS after 14-day of transplantation
Femurs
Mouse 1
Mouse 2
lung fragments
Ossicles
Mouse 1
Mouse 2
Ossicles
Femurs
Mouse 3
b
MDA-MB-231
DAPI Luciferase
c
Engraftment %
MDA-MB-231
d
PDX1
PDX2
e
CD56
PHOX2B
PDX1
PDX2
f
PDX1
Q5 60.3
Q6 32.0
Q8 6.85
Q7 0.91
GD2-BV786
CD56-BV605
PDX2
Q1 43.1
Q2 38.7
Q4 18.2
Q3 0.022
g
% out of GFP+ cells
GD2+
GD2+CD56+
PDX1-engrafted
PDX2-engrafted
h
PDX1
PDX2
Control

Fig.S4
