## Supplementary Figure Legends_Methods for "Engineering of human mini-bones for the standardized modeling of healthy hematopoiesis, leukemia and solid tumor metastasis"

**Fig.S1. MSOD-B cells selectively primed towards EC and IM ossification pathways can reproducibly form hOss. a-b,** Representative Safranin O **(a)** and Alizarin Red S **(b)** staining from *in vitro* Cho and Ost tissues after 3-week of differentiation. n = 3 biological replicates. **c-e,** Representative staining of MMP13 **(c)**, VEGF **(d)** and Laminin **(e)** of whole 100µm sections from Cho and Ost *in vivo* tissues explanted after 1-week of implantation. Nuclei are stained with DAPI. n = 3 biological replicates. Scale bar = 500µm. **f,** Number of MSOD-B (GFP) cells within 3-week differentiated *in vitro* and 1-week implanted Cho and Ost tissues. n = 6 biological replicates for *in vitro* samples and n = 3 biological replicates for *in vivo* samples. Statistical values were determined by two-way ANOVA followed by Šídák's post-hoc test, *p < 0.05, **p < 0.01. **g,** Representative 3D reconstruction of Col2 and ColX within 100µm thick Cho and Ost *in vitro* and *in vivo* tissues by Imaris software. Nuclei are stained with DAPI. n = 3 biological replicates. Scale bar = 500µm. **h,** Representative staining of OCN of whole 100µm sections from 3-week differentiated *in vitro* and 1-week implanted Ost tissues. Nuclei are stained with DAPI. n = 3 biological replicates. Scale bar = 500µm. **i,** Median fluorescence intensity (MFI) of OCN in 3-week differentiated *in vitro* and 1-week implanted Ost tissues. Shown are mean values + 1 SE. n = 3-6 biological replicates. Statistical values were determined by *t* test, ***p < 0.001.

**Fig.S2. MSOD-B hOss reconstitute a human mesenchymal niche supporting the establishment of human hematopoiesis. a-b,** Quantification of total volume (TV) **(a)** and bone volume (BV) **(b)** of Cho and Ost hOss explanted at 4, 12 and 24-week post-implantation. n = 3 biological replicates. Statistical values were determined by one-way ANOVA followed by Tukey post-hoc test, *p < 0.05, **p < 0.01, ***p < 0.001. **c,** Representative staining of human $CD45^+$ cells within 2 different Cho and Ost hOss explanted 24-week post-implantation. Nuclei are stained with DAPI. Scale bar = 500µm. **d,** Percentage of $hCD45^+$ cells in mouse peripheral blood (PB) at 8, 12, 16 and 20-week post-transplantation. n = 17 mice. **e,** Percentage of $CD3^+$ T cells, $CD19^+$ B cells and $CD33^+$ myeloid (My) cells out of $hCD45^+$ cells in PB at 8, 12, 16 and 20-week post-transplantation. n = 17 mice**. f,** Representative flow cytometry dot plots of MSOD-B-GFP cells in Cho and Ost hOss explanted at 29-week post-implantation. **g,** Representative macroscopic images of Cho and Ost hOss isolated at 7-week post-implantation. Scale bar = 0.5cm. **h,** Representative flow cytometry dot plots of MSOD-B-GFP cells in Cho and Ost hOss explanted at 7-week post-implantation. **i,** Frequency (%) of MSOD-B cells within Cho and Ost hOss at 7-week post-implantation. n = 3 biological replicates. Statistical values were determined by *t* test. *p < 0.05. **j,** Representative images of primary hOss-isolated and expanded MSOD-B cells at 7-week post-implantation. **k-l**, Representative flow cytometry histograms **(k)** and percentage **(l)** of CD73, CD90 and CD105 of hOss-isolated

MSOD-B cells at 7-week post-implantation. n = 3 biological replicates. **m,** Representative macroscopic images of primary and secondary hOss explanted at 4-week post-implantation. Size = 0.4-0.5cm. **n,** Representative histological images of Masson's trichrome staining of the sections from 4-week implanted secondary hOss. n = 3 biological replicates. BM (bone marrow), CB (cortical bone), TB (trabecular bone), V (vessels), blc (bone-lining cells), oc (osteocytes), ob (osteoblasts), a (adipocytes) shown with arrows.

**Fig.S3. Primary acute myeloid leukemia cells preferentially home and engraft in MSOD-B hOss. a,** Clinical information of AML patient samples. **b,** Percentage of $hCD45^+$ leukemic cells in PB and the spleen of NSG mice at 24-week post-intravenous (IV) and intra-ossicle (IO) transplantations. n = 2-4 NSG mice per AML-patient. **c,** Representative staining and quantification (%) of $hCD45^+$ (terracotta) AML cells within hOss at 24-week post-IO and IV transplantations. Mouse bone marrow (mBM) cells and MOLM-13 (AML cell line) cells were used to test antibody specificity. Scale bar = 50µm. **d,** Percentage of $CD19^+$ B and $CD3^+$ T cells, $CD13^+$, $CD33^+$, $CD34^+$, $CD34^+CD117^+$ cells out of $hCD45^+$ AML cells in patient (PAT) initial samples compared to hOss at 24-week post-IO transplantation. n = 2-4 biological replicates per AML patient sample, 8 AML-engrafted patient samples in total.

**Fig.S4. MSOD-B hOss support breast cancer metastasis and neuroblastoma cell engraftment. a,** Representative bioluminescence images showing the *ex vivo* distribution of MDA-MB-231 cells in hOss and mouse femurs at 14-day post-transplantation. **b,** Representative staining of Luciferase-MDA-MB-231 cells within hOss at 14-day post-transplantation. Nuclei are stained with DAPI. Scale bar = 50µm. **c,** Percentage of engrafted-hOss with MDA-MB-231 cells. n = 12 biological replicates. **d,** Representative CT/bioluminescence images showing the *in vivo* distribution of NB PDX1 and PDX2 cells within hOss. **e,** Representative staining of NB markers hCD56 and hPHOX2B in NB PDX1 and PDX2 cells within hOss. Scale bar=20µm. **f-g,** Representative flow cytometry plots **(f)** and frequency (%) **(g)** of $GD2^+$ and $GD2^+CD56^+$ cells in hOss. n = 3 biological replicates. **h,** Representative macroscopic images of control and NB PDX1 and PDX2-engrafted hOss. Size = 0.9-1cm for PDX1-engrafted hOss. Size = 0.4-0.5cm for PDX2-engrafted and control hOss.

### Methods

#### Mice

NOD.Cg-*Prkdc*$^{scid}$ *Il2rg*$^{tm1Wjl}$/SzJ (NSG) mice (2-4-month-old) were obtained from Charles River Laboratories or from an internal divisional stock. All mouse experiments and animal care were performed in compliance with the Lund University Animal Ethical Committee (15485–18, 19012-19) and according to protocols approved by the Danish Animal Ethical Committee (P21-

149) and in accordance to the 3R's principles. Mice were housed at a standard day-night cycle in an IVC-cage at a positive air pressure and were handled aseptically under a laminar flow. Mice were fed with autoclaved diet and water ad libitum. During the implantation procedure at BRIC institute in Copenhagen, mice were administered with Norodyl (Carprofen) (Scanvet) subcutaneously at 5 mg/kg and Bupaq Multidose (Buprenorphine) (Salfarm Scandinavia) at 0.1 mg/kg for the induction of analgesia. In addition, they received water for 4 days ad libitum supplemented with Carprofen at (0.067 mg/l). Temgesic was administered after implantation at Lund University. The anaesthesia was induced and maintained with 2-3 % Isoflurane (Attane) during the implantation, and to avoid heat loss the mice were kept on a heating pad during the procedure.

**Generation of MSOD-B hOss**

MSOD-B cells from passage 20-23, expressing hTERT-GFP, inducible caspase 9 (iC9)-ΔCD19 and BMP2-ΔNGFR [1], were seeded at a density of 3200 cells/cm$^2$ and cultured until reaching to 80-90% confluency in α-minimum essential medium (αMEM) supplemented with 10% fetal bovine serum (FBS), 1% HEPES (1M), 1% sodium pyruvate (100mM), 1% of penicillin-streptomycin-glutamine (100X) solution (all from Gibco) and 5 ng/ml of fibroblast growth factor-2 (FGF-2, R&D Systems), in a humidified 37 °C/5% $CO_2$ incubator. The medium was changed twice a week. Afterwards, cells were trypsinized with Trypsin-EDTA (0.25%) (Gibco) and used for generating *in vitro* tissues. 3.5 million MSOD-B cells were seeded onto type I collagen meshes (cylinder of 8mm in diameter, 3mm thick; Ultrafoam, Davol), corresponding to a density of 3.5 x 10$^3$ cells/cm$^3$. Tissues were cultured for 3 weeks in chondrogenic medium (DMEM supplemented with 1% penicillin-streptomycin-glutamine (Gibco), 1% HEPES (1M) (Gibco), 1% sodium pyruvate (100mM) (Gibco), 1% ITS (100x) (Insulin, Transferrin, Selenium) (Gibco), 0,47mg/ml linoleic acid (Sigma), 0.12% bovine serum albumin (BSA) (Sigma), 0.1mM ascorbic acid (Sigma), 10$^{-7}$M dexamethasone (Sigma) and 10ng/ml TGF-β3 (Novartis)) and in osteogenic medium (αMEM supplemented with 10 % FBS, 1% penicillin-streptomycin- glutamine (Gibco), 1% HEPES (1M) (Gibco), 1% sodium pyruvate (100mM) (Gibco), 0.1mM ascorbic acid (Sigma), 10$^{-4}$M dexamethasone (Sigma) and 0.01M beta-glycero-phosphate (Merck). After 3 weeks, MSOD-B-derived *in vitro* engineered tissues were subcutaneously implanted in NSG mice for the formation of hOss. The number of engineered tissues per animal varied depending on the type of transplantation: 4-6 implants for the CB-CD34$^+$ cell, 2 implants for AML patient cell and 4 implants for BC or NB PDX cell transplantations.

**Histological staining**

MSOD-B *in vitro* tissues or *in vivo* hOss were washed in 1XPBS and fixed overnight with 4%

formalin (Solveco AB), at 4°C. The hOss were decalcified with 10% EDTA, pH=8, at 4°C for 2 weeks before paraffin-embedding. Tissues were sequentially dehydrated in 35%, 70%, 95%, 99.5% graded ethanol solution (Solveco AB) twice for 20 minutes each. This was followed by 10min washing in 99.5% ethanol/xylene solution (1:1, Fisher Scientific) and then 20 min in xylene (Fisher Scientific), twice. After immersing tissues in paraffin at 56°C overnight, they were embedded and cut with a microtome in 7-10 µm sections. The sections were then dried overnight at 37°C. To remove the paraffin, sections were washed with xylene for 7min twice, then once in 99.5% ethanol/xylene solution (1:1) for 3min. Afterwards, sections were hydrated in 99.5%, 95%, 70%, 35% ethanol, twice for 7min each.

***Safranin O staining***

Sections were stained with Mayer's hematoxylin solution (Sigma-Aldrich) for 10min. This was followed by washing step with distilled water to remove extra color from sections. Then they were stained with 0.01% fast green solution (Fisher Scientific) for 5min, and extra color was removed by rinsing the sections quickly with 1% acetic acid solution (0.5 ml acetic acid in 49.5 ml distilled water) for 15 seconds. After staining the slides with 0.1% safranin O (Fisher Scientific) solution for 5min, dehydration and clearing were done by immersing the slides in 95% and 99.5% graded ethanol, afterwards with 99.5% ethanol/xylene solution (1:1). The sections were finally washed in xylene twice for 2min to remove ethanol followed by mounting with PERTEX mounting medium (PERTEX, HistoLab).

***Alizarin Red S staining***

The tissue sections were stained with 1% aqueous alizarin red S (PH=4.2, Sigma, A5533) for 1min. This was followed by clearing the slides by rinsing in 60% isopropanol (Sigma-Aldrich), and in absolute isopropanol for 30min each before washing in xylene and mounting using mounting medium (PERTEX, HistoLab).

***Masson's trichrome staining***

Trichrome staining was performed using Trichrome Staining Kit (Sigma-Aldrich Sweden AB) according to the guideline of the manufacturer. The tissue sections were deparaffinized and immersed in cold running deionized water for 3min. Then the sections were kept in Bouin's solution, at room temperature, overnight or at 56°C for 15min. The slides were washed by running tap water and stained using working Weigert's iron hematoxylin solution (equal volumes of solution A with B) for 5min for nuclei detection (in black). After washing the slides, the cytoplasm was stained in red with Biebrich Scarlet-Acid fuchsin for 5min followed by clearing the slides by immersing in working phosphotungstic/ phosphomolybdic acid solution (25ml P. tungstic + 25ml P. molybdic+ 50ml distilled water) for 5min. Collagen was stained in blue by immersing the slides in aniline blue solution for 5min followed by clearing in 1% acetic acid solution diluted in distilled water (glacial, Fisher Scientific) for 2min and washing with running deionized water. Finally, the sections were dehydrated in graded ethanol solutions

(95% once, 100% twice) for 2min each before washing with xylene twice for 2min and mounted with PERTEX mounting medium.

**Cell transplantation**

*Human cord blood CD34+ cells*

The human cord blood CD34+ (CB-CD34+) cells were isolated by using CD34 MicroBead Kit UltraPure, human (Miltenyi Biotec) according to the manufacturer protocol. Four-week post implantation, 1.5 x $10^5$ CB-CD34+ cells (pooled from a minimum of 3 donors) were injected intravenously into mice post 24h sub-lethal irradiation (200cGy). Peripheral blood (PB) chimerism was determined by flow cytometry every 4 weeks up to 20 weeks post-transplantation. The transplantation experiment was performed 2 times with a cohort of five recipient mice (each mouse bearing 4-6 hOss) per transplant. In general, transplanted mice were regarded as engrafted when BM chimerism was higher or equal to 1%.

*Human AML patient samples*

Frozen bone marrow aspirates from 10 AML patients were obtained with a written informed consent in accordance with the Declaration of Helsinki under the following ethical approval: Copenhagen; 1705391. The cells were thawed with RPMI1640 (Sigma) supplemented with 50 units of DNAse I (Sigma), 20% foetal bovine serum (FBS), 1% penicillin/streptomycin (Pen/Strep), 1 mM sodium pyruvate (Gibco) and 10mM HEPES (Gibco), the CD45+/CD3-/CD19-/CD34+ or CD45+/CD3-/CD19- populations were sorted with the FACS Aria III (BD Bioscience) using the following mouse anti-human monoclonal antibodies: CD45, V450 (clone HI30, BD, 1:100); CD3, APC-H7 (clone SK7, BD, 1:50); CD19, BV786 (clone HIB19, BD, 1:200), CD34, BV605 (clone 8G12, BD, 1:100) and FITC CD38 (clone HIT2, Biolegend, 1:50). The sorted cells were re-suspended with Myelocult H5100 (Stem cells tech) in a volume of 15 µl for intra-ossicle and 250 µl for intravenous (tail-vein) injections. Four-week post implantation the cells were injected with a 29 G syringe (BD Bioscience) into sub-lethally irradiated NSG mice (24h before transplantation, 200cGy for the mice bearing hOss and 250cGy for the mice not bearing hOss). The mice bearing hOss (2 hOss per mouse) were implanted as one patient in two mice. For $CD45^+/CD3^-/CD19^-$ cells, 250´000 cells were injected per ossicle (total 500´000 cells per mouse, n=2-4 mice per AML patient sample) and 500000 cells intravenously per mouse (n=2-4 mice bearing ossicles, n=1-2 not bearing ossicles per AML patient sample). For $CD45^+/CD3^-/CD19^-/CD34^+$ cells, 100´000 cells were injected per hOss (total 200´000cells per mouse) and 200000 cells intravenously per mouse (hOss and no hOss bearing). In general, transplanted mice were regarded as engrafted when BM chimerism was higher or equal to 1%.

*Breast cancer cell lines*

Four-week post implantation $10^6$ cells of the luciferase-transduced MDA-MB-231-LUC or the MCF7-LUC breast cancer cell lines were intravenously administered via tail-vein injection into 8-12 weeks-old NSG female mice (4 mice per cell line), 3 mice bearing 4 hOss per mouse or 1 mouse with no hOss as a control. Tumor engraftment was assessed *in vivo* via bioluminescence imaging.

*Human neuroblastoma PDX cells*

Neuroblastoma (NB) patient-derived xenograft (PDX) cells were grown as tumor organoids in serum-free medium as previously described [2, 3]. Cells from PDX models LU-NB-1 (PDX1) and LU-NB-2 (PDX2) were transduced with a luciferase and enhanced green fluorescent protein (eGFP) expressing third-generation lentiviral vector, constructed as previously described [4]. Four-week post-implantation, either NB PDX1 or PDX2-GFP-LUC cells (n=$10^6$ cells) were injected directly into 2 out of 4 hOss per mouse in five female NSG mice at a volume of 15µl. Additionally, control mice were intra-femorally injected (left femur) with $10^6$ NB PDX1 or PDX2 cells in a volume of 7µl. Tumor engraftment was assessed *in vivo* via bioluminescence imaging.

**Flow cytometry**

*CB-CD34+ cells*

Bone marrow (BM) cells from mouse bones and hOss were isolated by flushing mouse bones with 21-22G needles and by crashing ossicles with micro pestles in 1.5ml tubes using PBS/2% FBS/2mM EDTA. Mononuclear cells were isolated by low-density centrifugation (Histopaque 1077, Sigma) and stained according to the standard procedures. Samples were analyzed on a LSR Fortessa flow cytometer (BD Biosciences). DAPI (4', 6-Diamidino-2-Phenylindole, Dihydrochloride) was used to discriminate dead cells with the final concentration of 0.2ug/ml. Monoclonal antibodies to human CD45, Alexa Fluor 700 (clone HI30, BD) and mouse CD45, APC-Cy7 (clone 30-F11, BioLegend) were used to distinguish human from mouse cells. The antibodies used for PB and BM lineage analysis were all mouse anti-human: CD3, PE-Cy7 (clone UCHT1, BD), CD19, APC (clone HIB19, BD) and CD33, PE (clone WM53, BD). Lineage flow cytometry analysis data are plotted as the percentage of $CD19^+$ B cells, $CD3^+$ T cells and $CD33^+$ myeloid (My) cells among donor-derived cells. For early hematopoiesis analysis samples were stained with anti-human CD38, PE (clone HIT2, BD), anti-human CD34, APC (clone 581, BD), anti-human CD90, PE-Cy7 (clone 5E10, BD), anti-human 45RA, PerCP-Cy5.5 (clone HI100, BD), and anti-human hematopoietic lineage (Lin), FITC (eBioscience, 15558316). The flow cytometry analysis data were plotted as a percentage of hematopoietic stem cells (HSCs, gated as $CD38^-CD34^+CD90^+CD45RA^-$), multipotent progenitors (MPPs, gated as $CD38^-CD34^+CD90^-CD45RA^-$) and lymphoid-primed multipotent progenitors (LMPPs, gated as $CD38^-CD34^+CD90^-CD45RA^+$) distribution among donor-derived $CD45^+Lin^-$ cells. Samples were analyzed with FlowJo (version 10.7, BD Biosciences).

*AML patient samples*

The mice were euthanized after 24 weeks and the femoral bones, hOss, spleen and peripheral blood were collected. Bones were crushed and filtered. Red blood cell lysis was performed with ACK Lysis buffer (Thermo Fisher Scientific) on all samples except the hOss. The hOss were dissociated into a single cell suspension with a gentleMACS Dissociator (Miltenyi Biotech). A Nucleocounter NC-250 (Chemometec) was thereafter used to count the cells. The following markers were used to analyse the samples: The isolated single cells were screened for human CD45, V450 (clone HI30, BD, 560367, 1:100) content and myeloid cells with human CD33, PE (clone WM53, Thermo Fisher Scientific, 12-0338-43, 1:200), CD13, BV510 (clone L138, BD, 745015, 1:200), CD34, BV605 (clone 8G12, BD, 745247, 1:100), CD117, PE-Cy7 (clone 104D2, Thermo Fisher Scientific, 25-1178-42, 1:50) and for lymphoid cells; human CD3, BV650 (clone SK7, BD, 563999, 1:50) and CD19, BV786 (clone HIB19, BD, 740968, 1:200). The percentage of human CD45 was calculated from a 2D plot with both human CD45 and mouse CD45. Sequential Boolean plotting were then used to determine the other markers based on the percentage of human CD45 cells. The flow cytometric panel was as well applied on the primary cells to be able to compare the patient-derived xenograft (PDX) material; gating scheme is outlined in the supplementary material.

Acquisition of flow data was performed with DIVA (BD Bioscience) and a flow cytometer (Celesta, BD Bioscience), gating and compensation was performed with FlowJo v10.7.2 (BD Bioscience) and Prism v9.1 (GraphPad) was used for statistical evaluation. Study data were collected and managed using REDCap electronic data capture tools hosted at University of Copenhagen [5].

*NB PDX cells*

The hOss were explanted and processed by enzymatic digestion as described before [6]. Depending on the red blood cell content as determined by the color of the cell pellets, we performed an additional ficoll layering and/or a short RBC lysis step. Cells were blocked for 15 min at room temperature with human and murine Fc receptors (Human TruStain FcX and anti-mouse CD16/32, from BioLegend), and subsequently stained with the following antibodies: anti-mouse CD45, Pacific Blue (clone: 30-F11, BioLegend), anti-human CD56 (NCAM), Brilliant Violet 605 (clone HCD56, Nordic Biosite) and anti-human Disialoganglioside GD2 Brilliant Violet 786 (clone 14.G2a, BD Biosciences). Unstained cells were used to define forward and side scatter properties and 7AAD staining (Thermo Scientific) was used to discriminate living cells. Single-color tubes served to define the compensation matrix. Gates were set according to their corresponding fluorescence-minus-one (FMO) controls after doublets and dead cells exclusion as well as gating for CD45 negative populations. NB PDX1 and PDX2 populations were identified as positive for GFP and the combination of CD56 and GD2. Samples were run on a BD LSRFortessa or a BD LSRFortessa X20 flow cytometer (BD Biosciences) and analyzed with FlowJo (version 10.7, BD Biosciences).

*hOss-isolated MSOD-B cells*

The hOss were explanted at 7 and 29-week post-implantation and processed by enzymatic digestion using Liberase TM (Merck) (26 units/ml, working concentration: 0.26 unit/ ml) at 37°C for 45min. The samples were inactivated, homogenized and passed through 70 µm strainers to remove clumps. Depending on the red blood cell content as determined by the color of the cell pellets, additional ficoll layering and/or a short RBC lysis step was performed. Cells were blocked for 15 min at room temperature with Fc receptors (Human TruStain FcX, from BioLegend), and subsequently stained with anti-human NGFR, PE (clone: C40 1457, BD). For the isolation and expansion of MSOD-B cells, after removing red blood cells, $10^6$ cells were seeded and cultured in α-minimum essential medium (αMEM) supplemented with 10% fetal bovine serum (FBS), 1% HEPES (1M), 1% sodium pyruvate (100mM), 1% of penicillin-streptomycin-glutamine (100X) solution (all from Gibco) and 5 ng/ml of fibroblast growth factor-2 (FGF-2, R&D Systems), in a humidified 37 °C/5% CO2 incubator. After 1-2 weeks GFP+ MSOD-B cells were collected for the immunophenotypic profiling by using the following antibodies: anti-human CD105, APC (clone 266), anti-human CD73, PE (clone AD2), anti-human CD90, PE-Cy7 (clone 5E10), all from BD Biosciences.

**Microtomography (µCT)**

The µCT of the explants was performed with the U-CT system (MILABS, Netherland) using a tungsten x-ray source at 50kV and 0.21mA. For each sample, a circular scan (360°) was recorded with an incremental step size of 0.250°. Volumes were reconstituted at 10µm isotropic voxel size using MILABS software analysis. For bone volume analysis, the highly mineralized tissue volume was quantified using Seg3D (v2.2.1, NIH, NCRR, Science Computing and Imaging Institute (SCI), University of Utah, USA). For total volume analysis, each sample was meshed with Blender (v2.82a, Netherland) and analyzed with an in house developed script.

**Intracellular flow cytometry staining**

After 3 weeks, MSOD-B-derived *in vitro* engineered tissues and 1 week *in vivo* implanted tissues were digested with collagenase II (300 U/mg) (Thermo Scientific), Collagenase P (2 U/mg) (Sigma), 2mM Calcium Chloride (Sigma) in PBS for 60min, in a humidified 37 °C/5% $CO_2$ incubator. At the end of the digestion, cold α-minimum essential medium (αMEM) supplemented with 10% fetal bovine serum (FBS), 1% HEPES (1M), 1% sodium pyruvate (100mM) and 1% of penicillin-streptomycin-glutamine (100X) solution (all from Gibco) to stop the digestion. The samples were fixed and permeabilized with Cytofix/Cytoperm Solution (BD

Biosciences) according to the manufacturer protocol. Then the samples were incubated with 10% donkey serum (Sigma) in BD Perm/Wash Buffer (BD Biosciences) for 30 min. Primary and secondary antibody incubations were performed at RT in BD Perm/Wash Buffer (BD Biosciences) for 1h and 30min, respectively. The primary antibodies: rabbit anti-human Sox9 (Thermo Scientific, PA5-85275), rabbit anti-Collagen Type X (ColX) (abbexa, abx101469), rabbit anti-Collagen II (GeneTex, GTX20300), rabbit anti-Osteocalcin (Bioss, bs-4917R). The secondary antibody: donkey anti-rabbit DyLight649 (BioLegend).

**Immunofluorescence staining (IF)**

*In vitro* or *in vivo* samples were fixed in 4% paraformaldehyde (Thermo Scientific) at 4°C, for 24h and decalcified with 10% EDTA solution (Sigma), pH=8 up to 2 weeks with gentle rocking, at 4°C. Embedding was performed using 4% low-melting agarose (Sigma) and 100μm thick sections were cut using a 7000smz vibratome with stainless steel or ceramic blades (Campden). All steps of IF were done with gentle rocking. Sections were blocked and permeabilized with 0.5% Triton X-100 (Sigma) in PBS and 20% donkey serum (Jackson ImmunoResearch) for 1h, at room temperature (RT). After blocking/permeabilization, sections were stained with primary antibodies at 4°C, overnight, and highly cross-absorbed secondary antibodies for 2-3h at RT with 3 times 20min washes in between using 0.1% Triton X-100 in PBS and 2% donkey serum. Tissue sections were treated with Vector TrueView Autofluorescence Quenching Kit to remove unwanted fluorescence (Vector Laboratories). Tissue sections were mounted with a Vectashield antifade mounting medium with DAPI (Vector Laboratories, H1200). Samples were imaged with an LSM780 confocal microscope (Zeiss) with a 20x objective. Primary raw data were imported into the Imaris9.5 Software package for further processing and conversion into 3-dimensional images.

Primary antibodies: rabbit anti-Laminin (Novus Biologicals, NB300-144), goat anti-human Sox9 (R&D Systems, AF3075), mouse anti-Collagen II (Invitrogen, MA137493), rabbit anti-Collagen Type X (ColX) (abbexa, abx101469), rabbit anti-Osteocalcin (Bioss, bs-4917R), rabbit anti-MMP13 (Bioss, bs-0575R), rabbit anti-VEGF (Bioss, bs-0279R), chicken anti-GFP (Aves Labs, GFP-1020), goat anti-Perilipin1 (abcam, ab61682), mouse anti-human CD45 (eBioscience, 14-0459-82), mouse anti-human/canine NGFR/TNFRSF16 (R&D Systems, MAB367), rabbit anti-PHOX2B (abcam, ab183741), goat anti-Luciferase (Novus Biologicals, NB100-1677). Secondary antibodies: Alexa Fluor488 donkey anti-chicken (Jackson ImmunoResearch, 703-545-155), Cy3 donkey anti-goat (Jackson ImmunoResearch, 705-165-003), CF568 donkey anti-rabbit (Biotium, 20098), CF633 donkey anti-mouse (Sigma-Aldrich, SAB4600131).

**Bioluminescence imaging**

Establishment of tumor cells within the hOss was monitored via non-invasive bioluminescence imaging using IVIS-CT spectrum (PerkinElmer). The mice were anesthetized using 2-3% isoflurane and injected subcutaneously with D-luciferin (150mg/kg, PerkinElmer) in PBS. Mice were placed inside the imaging chamber onto a heated stage under continuous exposure to isoflurane. For 2D imaging, the exposure was set to the maximum of 300s and sequential images were taken with 5 min intervals for 20 min. When a strong signal was detected in 2D, an additional 3D µCT scan was performed to increase the visualization of hOss and tumor cell position. Images were subsequently analyzed with Living Image 4.5.5 software (PerkinElmer). Regions of interest were selected and quantified as photons per second following deduction of the average background signal.

**Immunohistochemistry (IHC-P)**

*AML-engrafted hOss*

Decalcified (as mentioned above) hOss from four different PDX mice were 4% formalin-fixed, paraffin-embedded (FFPE) and sectioned at 4µm. The sections were then stained with Mayer's Haematoxylin and Eosin (Sigma); Masson's Trichrome-Goldner (Sigma); and with rabbit anti-human CD45 (clone EP322Y, Abcam, ab40763) and Horseradish Peroxidase (HRP) (Sigma); and were as well fluorescently labelled with rabbit anti-human CD45 (clone EP322Y, Abcam, ab40763, 1:250), chicken anti-GFP (Aves Lab, GFP-1020, 1:1000), 4′,6-diamidino-2-phenylindole (DAPI) and as well with rat anti-mouse CD45 (clone 30-F11, Thermo Fisher Scientific, 14-0451-85, 1:200) for technical evaluation on one ossicle and on MOLM-13 cells and normal NSG mouse bone marrow. The following secondary antibodies from Thermo Fisher Scientific were used: goat anti-rabbit AF555Plus (A32732, 1:1000) and goat anti-rat AF647Plus (A-21247, 1:1000). The material was formalin-fixed, heat-induced epitope retrieval (HIER) was performed with 10 mM citrate buffer (pH 6.0) at 96 °C for 15 min. Blocking of non-specific proteins was then performed with 10 % normal goat serum (Thermo Fisher Scientific, 31873/11819220) and 1% bovine serum albumin (Sigma). Acquisition of the IHC-P slides was performed at 40x with NanoZoomer (Hamamatsu) and the whole fluorescently labelled sections were imaged at 40x with tile scanning with an LSM800 confocal microscope (Zeiss). The images were analysed with either NDP.view2 v2.8.24 (Hamamatsu) or Zeiss Black/Blue v2.3/3.3 (Zeiss).

*Neuroblastoma cell engrafted hOss*

The hOss were fixed in 4% paraformaldehyde overnight before decalcification in 10% EDTA-solution (pH 8). EDTA solutions were changed every 2-3 days and the decalcification continued for approximately 14 days until the bone tissue became soft. The hOss were then embedded in paraffin and cut in 4µm or 12µm sections. H&E (HistoLab Products AB) staining was used for assessment of histopathology. Other stainings were performed using Autostainer Plus (DAKO). Primary antibodies were NCAM/CD56 (1:50; NCL-L-CD56-501, Leica Biosystems), PHOX2B (1:1000; EPR14423, Abcam). Stained tissue sections were captured using an AxioScan slidescanner (Zeiss).

**Tartrate-Resistant Acid Phosphatase (TRAP) staining**

TRAP staining was performed in paraffin-embedded hOss tissue sections to assess the osteoclastic activity in the explanted hOss based on a modified method originally described by Burstone [7]. Briefly, deparaffinized slides were immersed in 0.2M acetate buffer at pH5.0 containing sodium acetate trihydrate and sodium tartrate (both from Sigma) for 20min. The slides were then incubated in fresh, pre-warmed staining solution containing 1 mg/ml Naphthol AS-MX phosphate powder and 1 mg/ml Fast Red TR Salt 1,5-naphthalenedisulfonate salt (both from Sigma) at 37°C for approximately 2h. Upon visual confirmation of bright red color development, the slides were rinsed and subsequently counterstained with Mayer's Hematoxylin solution (Sigma) for 1 min. Pictures of whole-tissue sections at 2.5x magnification were captured on a Zeiss Axiophot microscope and quantified using ImageJ (version 1.53j). In brief, after adjusting the scale, the pictures were converted to RGB stacks. Outlines of the whole tissue area for each ossicle were defined using the polygon tool and measured. The area within each selected ossicle outline corresponding to TRAP-positive osteoclasts was measured after adjusting the threshold accordingly. Ratios of positive TRAP signal versus the whole tissue area in each ossicle indicated the level of osteoclastic activity in tumor-injected and non-injected ossicles.

**Statistical analysis**

Data are plotted as mean + 1 standard error (SEM) unless differently stated. The SEM is used to indicate the precision of an estimated mean. Statistical analyses were determined with GraphPad Prism 8.4.2 or Prism v9.1. The statistical tests and *P* values are indicated in related figure legends. *P* values of <0.05 were considered significant. Each experiment was performed with a minimum of three biological replicates representing the number of engineered tissues/hOss. Exact number of hOss are mentioned in related figure legends. In transplantation experiments, samples were included in the analysis when engraftment was more or equal to 1% for cord blood CD34+ cells after 20-week post-transplantation and for AML patient samples after 24-week post-transplantation. Mice showing signs of sickness and/or showing signs of major disease involving hematopoietic and non-hematopoietic tissues were excluded from the analysis. Mice for the experiments were randomly chosen from suppliers.

### References


1. Pigeot, S. et al. Manufacturing of Human Tissues as off-the-Shelf Grafts Programmed to Induce Regeneration. *Adv Mater*, e2103737 (2021).
2. Braekeveldt, N. et al. Neuroblastoma patient-derived orthotopic xenografts retain metastatic patterns and geno- and phenotypes of patient tumours. *International journal of cancer* **136**, E252-261 (2015).

3. Persson, C.U. et al. Neuroblastoma patient-derived xenograft cells cultured in stem-cell promoting medium retain tumorigenic and metastatic capacities but differentiate in serum. *Scientific reports* **7**, 10274 (2017).
4. Hansson, K. et al. Therapeutic targeting of KSP in preclinical models of high-risk neuroblastoma. *Sci Transl Med* **12** (2020).
5. Harris, P.A. et al. The REDCap consortium: Building an international community of software platform partners. *J Biomed Inform* **95**, 103208 (2019).
6. Gulati, G.S. et al. Isolation and functional assessment of mouse skeletal stem cell lineage. *Nature protocols* **13**, 1294-1309 (2018).
7. Burstone, M.S. Histochemical demonstration of acid phosphatases with naphthol AS-phosphates. *J Natl Cancer Inst* **21**, 523-539 (1958).
